# Profilin-1 Deficiency Activates STING to Drive T Cell-Mediated Anti-Tumor Immunity in Breast Cancer

**DOI:** 10.64898/2026.06.05.730362

**Authors:** Ian Eder, Masoumeh Baghaei, Sudeep Maurya, Virginia Yu, Elijah Wilson, Abrahim Kashkoush, Jia-Jin Liu, Silvia Liu, Jianhua Luo, Walter J. Storkus, Partha Roy

**Author notes:** <u>Corresponding author:</u> **Partha Roy**, PhD, 306 CNBIO, 300 Technology Drive, University of Pittsburgh, Pittsburgh, PA 15219, USA.

## Abstract

Dysregulation of actin-binding protein Profilin1 (Pfn1) in tumor cells has prominent impacts on the tumor-intrinsic aspects of tumor progression. However, whether and how modulation of Pfn1 expression in tumor cells influences immune surveillance in cancer is not known. We utilized an inducible CRISPR/Cas9 knockout (KO) model to first demonstrate that triggering Pfn1 depletion in breast cancer cells leads to features of genomic instability (polyploidy, micronuclei, and DNA damage) and intrinsic defects in both homologous-recombination- and non-homologous end-joining-mediated double-stranded DNA repair. Pfn1-deficient breast cancer cells exhibit nuclear envelope abnormality and the accumulation of cytosolic DNA. This leads to activation of the nucleic acid-sensing cGAS-STING pathway and the type-I interferon (IFN) response including STING-mediated upregulation of pro-inflammatory chemokines. In an immunocompetent mouse model of breast cancer, triggering Pfn1 loss selectively in tumor cells promotes an immunogenic tumor microenvironment marked by a striking increase in intratumoral presence of CD8⁺ T cells, leading to a robust tumor regression. Pfn1 knockout-induced tumor regression requires an intact immune system and can also be reversed by CD8^+^ T cell depletion. Based on these findings, we conclude that Pfn1 loss in tumor cells leverages a type I IFN response to drive a T-cell-mediated anti-tumor response in breast cancer. These findings for the first time reveal promising therapeutic opportunities in targeting Pfn1-driven pathways to enhance immunotherapeutic outcomes in breast cancer.

**Significance Statement:** Expression of actin-binding protein Profilin-1 is frequently altered in cancer; yet how these changes impact the immune response against tumors is unclear. Here we show that triggering Profilin-1 depletion in breast cancer cells promotes features of genomic instability, defects in DNA repair, and cytosolic release of DNA. This activates the cGAS-STING pathway, triggering a type I interferon response and immune-cell-attracting signals that drive a CD8+ T cell-mediated anti-tumor immune response and tumor regression *in vivo*. Therefore, Profilin-1 could be a novel actionable target for achieving immunological benefit in breast cancer. On a broader level, our studies establish a conceptual framework of how dysregulation of actin cytoskeletal proteins can harness nuclear damage-sensing signaling to augment anti-tumor immune response in cancer.

## INTRODUCTION

Profilins (Pfns) belong to a family of small actin monomer-binding proteins that have well-established roles in regulating actin cytoskeletal dynamics in cells (1). Pfn1 is the most abundant and the only ubiquitously expressed form of Pfn in cells. The most prominent consequence of loss-of-function of Pfn1 is a reduced level of actin polymerization in cells (2, 3). In physiological contexts, this leads to defects in actin-driven biological processes, such as cell migration and proliferation (particularly in cytokinesis) (1, 4–6). Pfn1 is also involved in other facets of cellular functions including regulation of microtubule dynamics (7), lipid metabolism (8, 9), and mitochondrial activity (10).

Pfn1 expression is dysregulated in tumor cells in a variety of human adenocarcinomas but in a context-specific manner. While in some cancers (breast, pancreatic, advanced hepatocellular carcinoma) Pfn1 expression is reduced in tumor cells vs their normal counterparts (11–15), its expression is elevated in gastric and metastatic non-small cell lung cancer (16, 17). In clear cell renal cell carcinoma (ccRCC), however, this tumor cell-centric view is complicated by the fact that Pfn1 expression is predominantly elevated in the stromal compartment (most notably in tumor-associated vascular endothelial cells) of the tumor microenvironment (TME) (18, 19). To date, loss-of-function mutations of Pfn1 in cancer have been exclusively reported in osteosarcoma (20, 21). In almost all cancer contexts described above, modulating Pfn1 expression has been shown to cause prominent changes in the tumor-intrinsic aspects of tumor progression. Specifically, in the context of breast cancer, we previously showed that loss of Pfn1 expression has contrasting effects on early vs late steps of tumor progression (11). While reduced Pfn1 expression promotes motility/invasion of tumor cells (contrasting the pro-migratory role of Pfn1 in physiological contexts) enhancing their disseminative ability, Pfn1 appears to be essential for tumor-initiating ability and metastatic colonization of breast cancer cells (5, 11). On the other hand, overexpression of Pfn1 suppresses tumor growth, migration/invasion, and metastatic colonization of breast cancer cells, therefore suggesting that a balanced level of Pfn1 is optimal for tumor progression in breast cancer (5, 22, 23).

How tumor-extrinsic aspects of tumor progression are impacted by modulation of Pfn1 expression in cancer cells remains a critical gap in the literature. Recently, several independent studies involving diverse cell types (osteoblasts, pagetic osteosarcoma cells, keratinocytes, podocytes, microglia) support the premise that Pfn1 loss leads to various nuclear abnormalities (micronuclei, polyploidy, and DNA damage) and/or to genomic arrangements (copy number variation and chromothripsis) in affected cells (20, 24–26). However, the potential implications of these findings in disease contexts are unclear. While genomic instability accelerates the accumulation of mutations thereby promoting tumor growth, it can also create vulnerabilities of cancer cells. Specifically, genomic instability can promote T-cell mediated anti-tumor immunity by increasing tumor-intrinsic immunogenicity as well as fostering a tumor microenvironment (TME) conducive to the recruitment and sustained function of anti-tumor T cells (27, 28). In the present study, we provide the first proof-of-concept that loss-of-function of Pfn1 in tumor cells can harness nuclear abnormality-triggered intracellular signaling to modulate the anti-tumor immune response in breast cancer.

## RESULTS

### Pfn1 loss leads to nuclear abnormalities in breast cancer cells

To examine the effects of acute Pfn1 loss in breast cancer cells, we engineered GFP/luciferase-expressing sublines of 4T1 (a murine breast cancer cell line) and MDA-MB-231 (MDA-231; a human breast cancer cell line) cells for doxycycline (dox)-inducible CRISPR-Cas9-based knockout (KO) of the *Pfn1* gene. Immunoblot analysis confirmed near-complete Pfn1 depletion in both cell lines after five days of dox induction (**Figure 1A**). A distinguishing phenotype of Pfn1 loss in both cell lines was the induction of polyploidy, a characteristic feature of mitotic defects. However, the polyploidy severity (judged by the average number of nuclei per cell) was more drastic in 4T1 cells than in MDA-231 cells (**Figures 1B-C**), suggesting context-dependent cellular sensitivity to Pfn1 loss. In addition to polyploidy, Pfn1-deficient cells also exhibited an increased frequency of micronuclei (small extra-nuclear bodies containing DNA) formation. Notably, a greater proportion of micronuclei in Pfn1 KO cells also completely lacked lamin-B1 in their envelopes (**Figures 1D-E**), a feature of rupture-prone micronuclei (29). Micronuclei result from various nuclear abnormalities including formation of lagging chromosomes due to mitotic defects, unrepaired DNA double-stranded breaks (DSBs), improper DNA replication, and nuclear envelope defects (30, 31). Although the overall contents of both lamin-B and -A/C at the nuclear lamina were enhanced in Pfn1 KO cells, these cells exhibited features that are indicators of nuclear envelope instability, such as reduced nuclear circularity (suggestive of increased nuclear deformability) and increased nuclear membrane invagination (32) (**Figures 1F-I, S1**).

**Figure 1.**
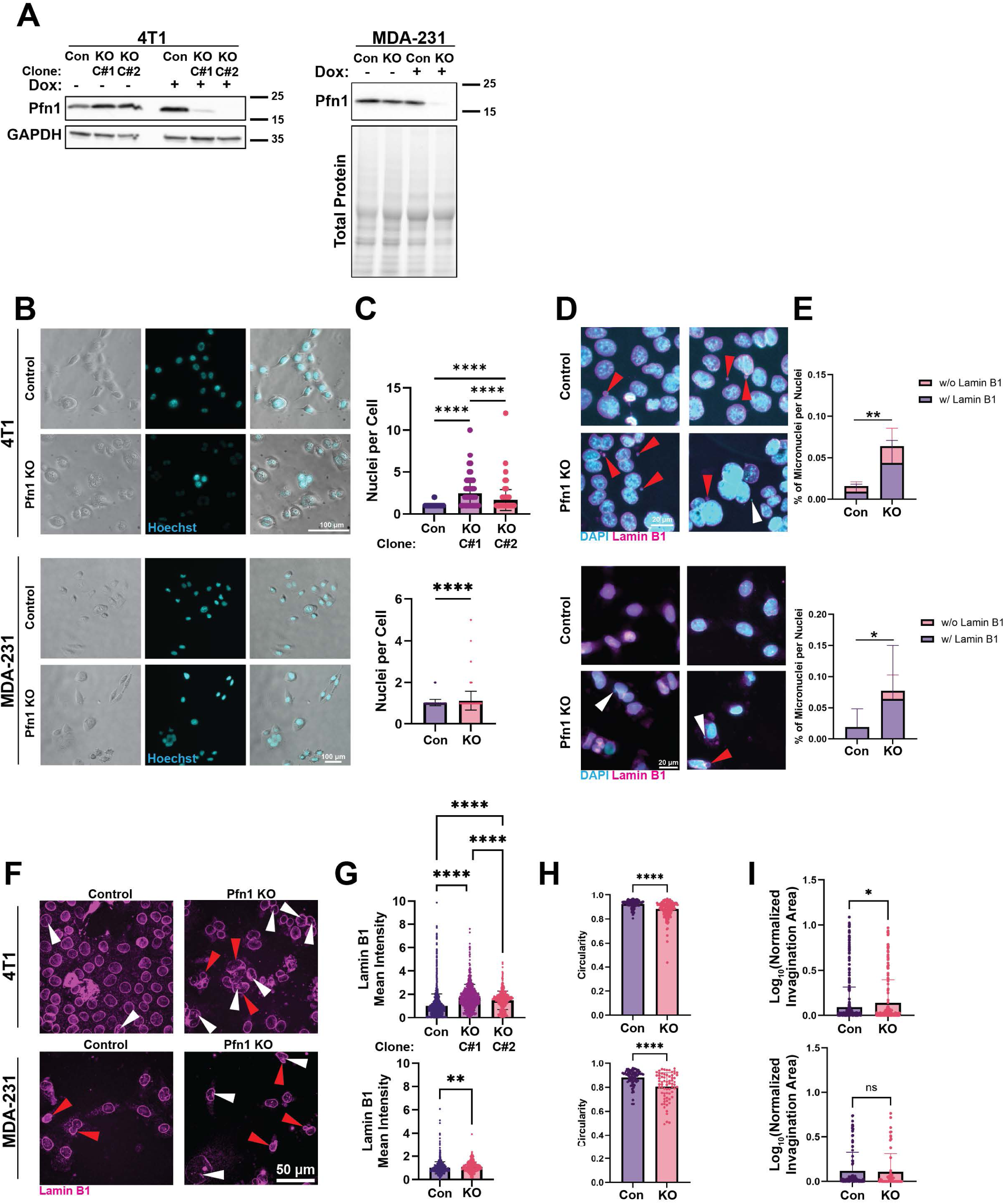
Pfn1 loss leads to nuclear abnormalities in breast cancer Cells. **A)** Pfn1 immunoblots demonstrating Cas9-mediated KO of the *Pfn1* gene in 4T1 cells and MDA-231 after 5 days of dox-administration. For 4T1 cells, 2 independent Pfn1 KO clones (C#1 and C#2) were analyzed. **B)** Live-cell phase contrast and Hoechst images of the indicated sublines (for 4T1, the image displayed in Pfn1 KO group represents clone C#1). **C)** Quantification showing the number of nuclei per cell in the indicated sublines. Data pooled from three independent experiments (4T1: Con = 474 cells, Pfn1 KO (C#1) = 307 cells, Pfn1 KO (C#2) = 229 cells; MDA-231: Con = 567 cells, Pfn1 KO = 566 cells). One-way ANOVA with a post-hoc Tukey test was used to compare groups. **D)** Representative immunofluorescence images of Lamin B1 and Hoechst-stained cells with micronuclei. Red arrows point to micronuclei with Lamin B1 staining. White arrows point to micronuclei without Lamin B1 staining (for 4T1, the image displayed in Pfn1 KO group represents clone C#2). **E)** Quantification of the number of micronuclei per cell and proportion of micronuclei with Lamin B1 staining. Data pooled from three (MDA-231) or two (4T1) independent experiments. (4T1: Con = 7 fields, Pfn1 KO (C#2) = 6 fields; 231: Con = 9 fields, Pfn1 KO = 9 fields). A Mann-Whitney Test was used to compare between groups. **F)** Representative immunofluorescence images of Lamin B1-stained 4T1 and MDA-231 cells. Red arrows point to nuclear invaginations. White arrows point to cells with low circularity (for 4T1, the image displayed in Pfn1 KO group represents clone C#2). **G)** Quantification of the mean intensity of Lamin B1 staining in the nuclear lamina region. Data pooled from three independent experiments. A student’s T test (MDA-231) or One-way ANOVA (4T1) with post-hoc Tukey’s test was used to compare between groups. (4T1: Con = 2301 cells, Pfn1 KO (C#1) = 934 cells, Pfn1 KO (C#2) = 572 cells; MDA-231: Con = 866 cells, Pfn1 KO = 488 cells). **H, I)** Quantification of the number of nuclear invaginations per cell and nuclear circularity. Data pooled from two independent experiments. A student’s T test was used to compare between groups. Number of cells analyzed for the invagination analysis: 4T1 (control = 493; Pfn1 KO (clone C#2) =190), and MDA-231 (Con = 98; Pfn1 KO = 63). Number of cells analyzed for the circularity analysis: 4T1 (Con = 109, Pfn1 KO (C#2) = 164) and MDA-231 (Con = 78; Pfn1 KO = 68). **** p < 0.0001, *** p < 0.001, ** p < 0.01, * p < 0.05.

In both cell lines, Pfn1 deficiency resulted in increased DNA damage marked by increased phosphorylation of H2A Histone family member X (H2AX) protein at Ser139 (γ-H2AX – a post-translational modification of H2AX that occurs rapidly at the sites of DNA DSBs) (33) (**Figures 2A-B, Figure S2**). We utilized 4T1 cells to further examine whether DNA DSB repair mechanisms are impacted by loss of Pfn1. First, we performed immunofluorescence analyses of Rad51 and 53bp1, key components for HR (homologous recombination)- and NHEJ (non-homologous end-joining)-mediated DSB repair, respectively (34), which revealed increased nuclear foci of both Rad51- and 53bp-foci in 4T1 cells resulting from Pfn1 depletion (**Figures 2C-F)**. Next, to determine whether the intrinsic HR and NHEJ capabilities are affected by loss of Pfn1, we performed HR and NHEJ reporter biosensor experiments (35, 36). As schematically illustrated in **Figure S3A**, in these biosensor assays, cells harbor integrated reporter constructs containing a disrupted GFP gene. A site-specific endonuclease generates a defined DSB within the reporter. Successful repair via either HR or NHEJ restores functional GFP expression, which can be quantified by flow cytometry-based assessment of GFP+ cells. To use these biosensors, we generated additional polyclonal Pfn1 KO variants of 4T1 cell line that did not express GFP (**Figure S3B**). Using this approach, we observed a significant decrease in GFP+ positive cells in Pfn1 KO vs control 4T1 cells in both HR and NHEJ biosensor assays (**Figures 2G-H**). We were also able to confirm the HR findings in normal HEK-293 cells in a setting of siRNA-mediated transient knockdown of Pfn1 expression (**Figures S3C-E**). This reduced capacity to repair DNA double-strand breaks was mirrored in gene set enrichment analysis of our transcriptomic data of 4T1 cells, which showed that DNA double-strand break processing was enriched in control cells relative to Pfn1 KO variants (**Figure 2I**). Multiple chromatin remodeling factors (SMARCAD1, KAT2, and SETMAR) responsible for increasing double stranded DNA break accessibility (37–39), End resection nucleases (RBBP8, DNA2, and EXD2) essential for HR initiation (40–42), and RAD52 which aids in the recruitment of RAD51 (43) were transcriptionally attenuated in Pfn1 KO cells (**Figure 2J**). These data demonstrate that loss of Pfn1 expression leads to diminished HR and NHEJ capabilities resulting in DNA DSB accumulation in breast cancer cells.

**Figure 2.**
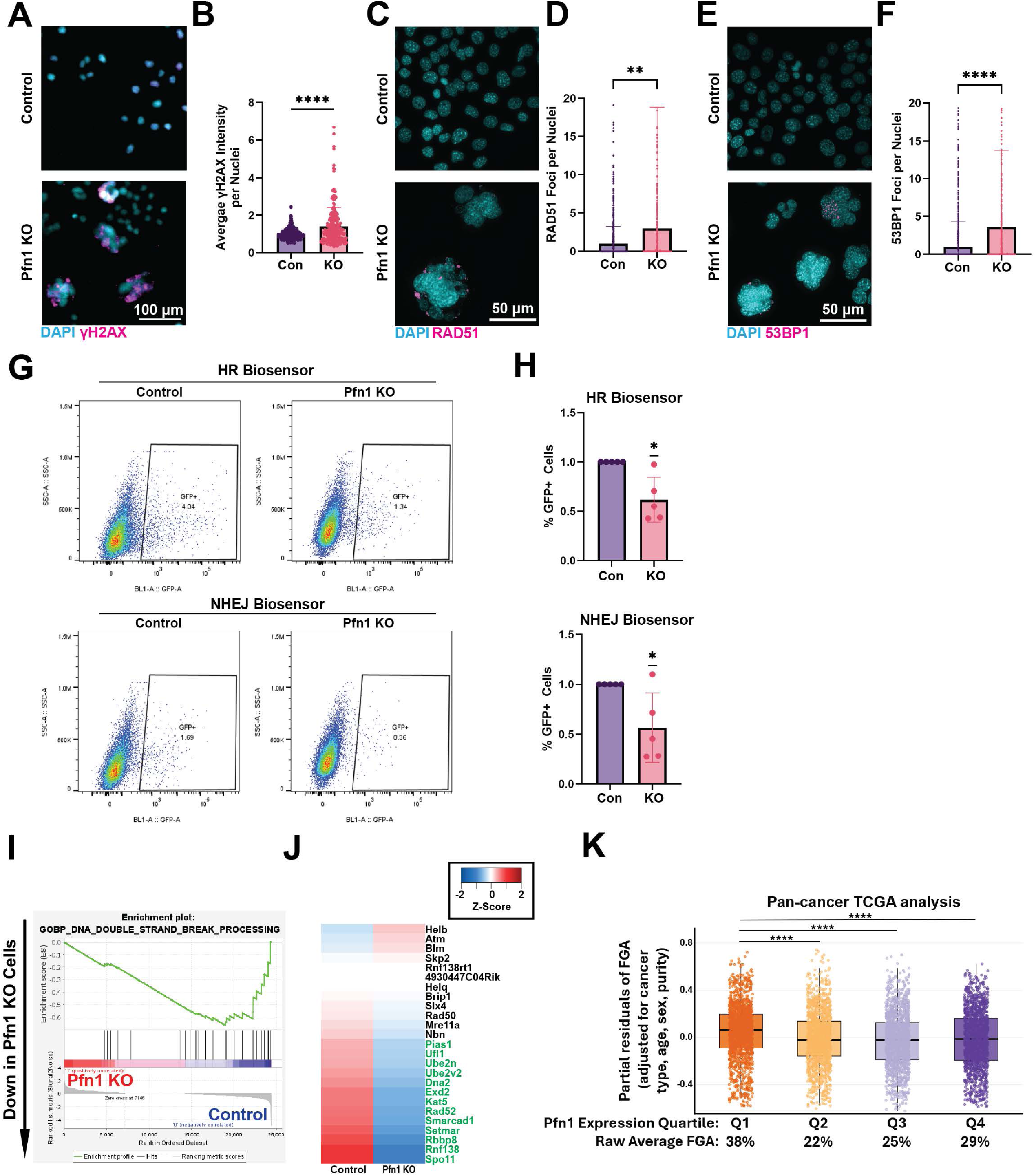
Pfn1 loss causes defects in DNA repair. **A, B)** Representative images and quantification of the mean intensity of γH2AX in the nucleus of control vs Pfn1 KO 4T1 cells. A student’s T test was used to compare between the two groups. Data pooled from three independent experiments. (Con = 553 cells, Pfn1 KO (C#2) = 241 cells). **C-F)** Representative images and quantification of RAD51(**C, D**) and 53BP1(**E, F**) foci per nuclei normalized to the nuclear area in control vs Pfn1 KO 4T1 cells. A student’s T test was used to compare between groups. Data pooled from three independent experiments. Number of cells analyzed for RAD51-foci analysis: Con = 1178; Pfn1 KO (C#2) = 668. Number of cells analyzed for 53bp1-foci analysis: Con = 1704; Pfn1 KO (C#2) = 635 cells. **G)** Representative flow cytometric data showing the proportion of control and Pfn1 KO 4T1 cells utilizing HR or NHEJ (GFP^+^) reporter assays. **H)** Quantification of the normalized proportion of control and Pfn1 KO 4T1 cells utilizing HR or NHEJ from five independent experiments. A one-sample T-test was used to compare experimental values to a hypothetical mean of 1. **I)** GSEA enrichment plots for the mouse GOBP_DNA_DOUBLE_STRAND_BREAK_PROCESSING gene set. Nominal p value = <0.0001, Normalized Enrichment Score = -1.32. **J)** Heat-plots showing Pfn1-dependent changes in the mRNA expression (represented as Z-scores) of genes in the GOBP_DNA_DOUBLE_STRAND_BREAK_PROCESSING. Genes shown in green text contributed to pathway enrichment (clone C#2 represented the Pfn1 KO group in the transcriptomic analyses of 4T1 cells). **K)** Clinical cancer exome sequencing data (TCGA) showing the fraction of the genome altered corrected for the specified covariates in samples stratified by Pfn1 expression compared to normal tissue controls. The raw average FGA shown below the graph refers to the average FGA of the indicated Pfn1 expression quartile prior to covariate adjustment. N = 10968 samples. A one-way ANOVA with post-hoc Tukey’s test. was used to compare between groups. **** p < 0.0001, *** p < 0.001, ** p < 0.01, * p < 0.05.

DSBs are the most deleterious forms of DNA damage, and inability to accurately repair DSBs lead to genomic instability and rearrangements. Accordingly, our pan-cancer analyses of The Cancer Genome Atlas (TCGA) dataset revealed highest fraction genome altered (FGA) in tumors associated with the lowest quartile of Pfn1 mRNA expression in human cancer (**Figure 2K**), further adding relevance to our findings in a human disease context. Collectively, these findings demonstrate induction of prominent nuclear abnormalities and DNA damage triggered by Pfn1 loss in cancer cells.

### Pfn1 deficiency triggers cytoplasmic accumulation of DNA and cGAS-STING activation

Since structurally unstable micronuclei and/or unrepaired DNA damage can lead to leakage of nuclear DNA into the cytoplasm (27, 44), we next quantified cytosolic DNA content in control vs Pfn1 KO cells using a qPCR-based detection method (44). In these experiments, nuclear and mitochondrial DNA were further distinguished by amplifications of 18S ribosomal DNA (rDNA) and mitochondrial cytochrome oxidase (COX) gene (mouse cells – Cox III; human cells - Cox II), respectively. In both 4T1 and MDA-231 cells, we detected increased cytosolic presence of nuclear DNA in Pfn1 KO group. While the cytosolic content of mitochondrial DNA was not significantly different between control and Pfn1 KO 4T1 cells, MDA-231 cells exhibited statistically significant increase in the cytosolic content of mitochondrial DNA in response to Pfn1 depletion (**Figures 3A**). This increased cytosolic DNA accumulation is unlikely to be explained solely by reduced TREX1-mediated DNA clearance, as expression of this major cytosolic exonuclease was not diminished in Pfn1 KO cells (**Figure 3B**). Thus, Pfn1 deficiency appears to promote cytosolic DNA accumulation through nuclear mechanisms in both breast cancer cell models, with an additional mitochondrial contribution in a cell-line-dependent manner.

**Figure 3.**
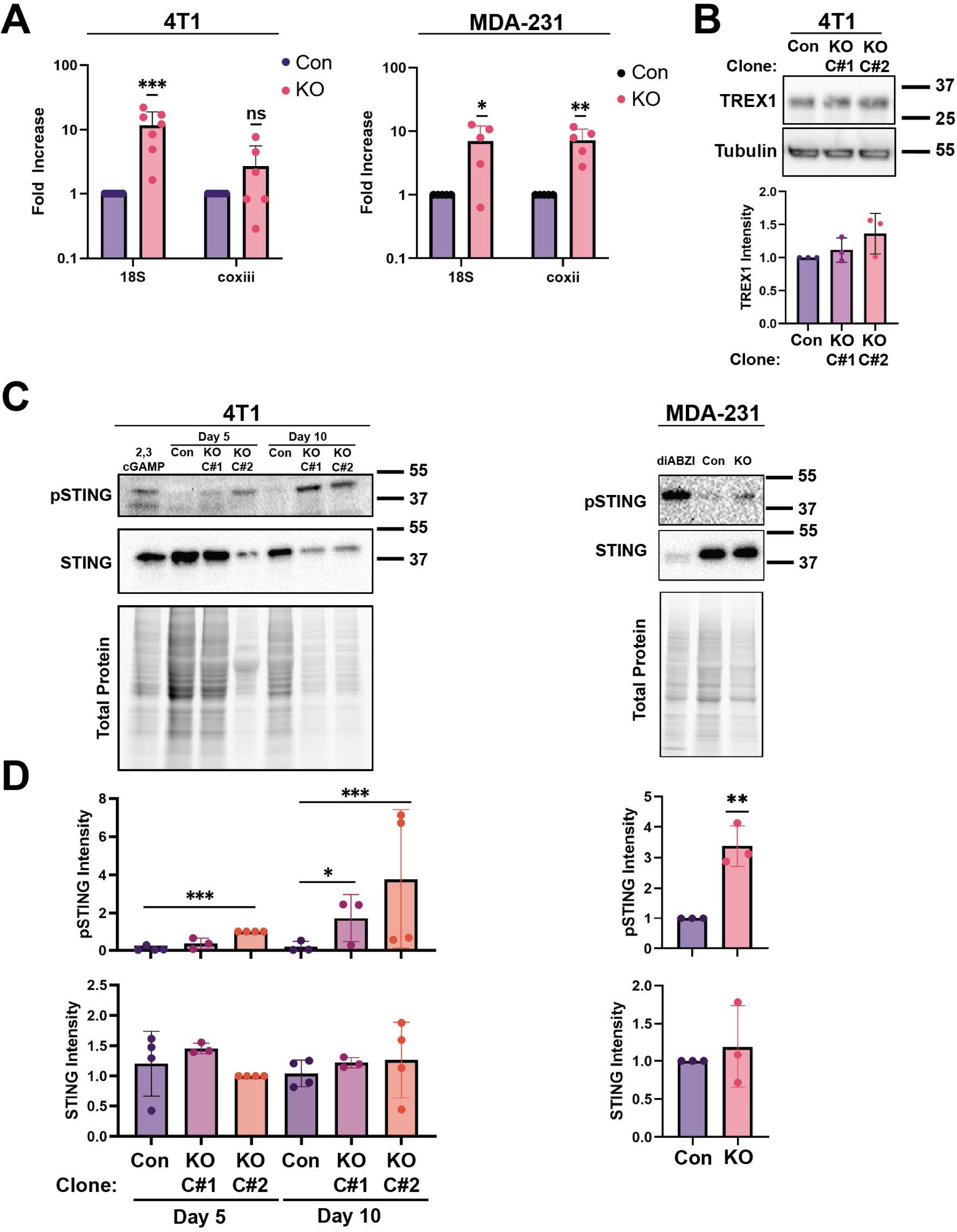
Pfn1 loss leads to cytosolic DNA accumulation driving STING activation. **A)** Quantification of relative levels of 18S and Coxiii (for 4T1 cells) and COXII (for MDA-231 cells) cytosolic DNA presence. A one-sample T-test was used to compare experiment values to a hypothetical mean of 1. The Pfn1 KO group for 4T1 cells represents clone C#2. **B)** Representative immunoblots and the associated quantification showing the relative expression levels TREX1 in the indicated cell lines. **C, D)** Representative immunoblots and the associated quantification showing the relative expression levels STING and pSTING (Ser365) in the indicated cell lines. 2’,3’-cGAMP transfection serves as the positive control for 4T1 cells and diABZI treatment serves as the positive control for MDA-231 cells. For STING and pSTING blots in MDA-231 cells, a one sample ratio T test was used to compare the experimental group to a hypothetical value of 1. For TREX1, STING, and pSTING blots in 4T1 cells, a mixed-effects REML model was used to compare values, with experimental replicate included as a matching factor to account for inter-experimental variability. Data were assumed to be lognormally distributed. Post-hoc Pairwise comparisons were assessed using a Tukey’s test. Only select significant comparisons are shown. **** p < 0.0001, *** p < 0.001, ** p < 0.01, * p < 0.05.

Cytosolic presence of DNA is a key upstream trigger for cGAS-STING activation (45). A key signature of cGAS-STING pathway activation is TBK1 (Tank-binding kinase 1)-mediated phosphorylation of STING upon cGAMP (cyclic guanosine monophosphate-adenosine monophosphate)-triggered dimerization of STING. Immunoblot analyses revealed an increase in STING phosphorylation in both 4T1 and MDA-231 cells five days after induction of Pfn1 KO (**Figures 3C-D**). STING activation by Pfn1 loss was also confirmed in a second Pfn1 KO 4T1 subline with a different Pfn1-targeting gRNA, further establishing the specificity of our findings (**Figure S4A**). We also noted that STING activation in 4T1 cells persisted even 10 days after triggering Pfn1 KO suggesting that this is a sustained cellular response (**Figures 3C-D**). Further, we observed increased nuclear localization of IRF3, a transcription factor which is directly phosphorylated by TBK1, after Pfn1 KO triggering (**Figures S4B-C**). Consistent with cellular senescence being frequently associated with chronic STING activation, we also observed a prominent senescent (but not apoptotic) phenotype in Pfn1 KO 4T1 cells (**Figure S5A-B**).

### Pfn1 loss triggers type I Interferon response and STING-dependent upregulation of pro-inflammatory chemokines

A key downstream consequence of cGAS-STING activation is a type I interferon (IFN) response, which is marked by increased production of type I IFNs, activation of JAK/STAT signaling, and STAT (primarily involving the action of STAT1/STAT2 heterodimers)-mediated upregulation of IFN-stimulated genes (ISGs) (46). Transcriptomic profiling of both breast cancer cell lines revealed enrichment of the type I IFN response (**Figure 4A –** significantly enriched in 4T1 cells and a trend towards enrichment in MDA-231 cells) and upregulation of ISGs (**Figure 4B**) in response to Pfn1 loss. For validation, we examined the expressions of 2′–5′-oligoadenylate synthetases OAS1 and OAS3 at the protein levels. OAS1 and OAS3 are canonical ISGs that synthesize 2′–5′-linked oligoadenylates in response to IFN signaling; these oligoadenylates activate RNase L to degrade viral and cellular RNA as part of the antiviral response (47). Immunoblot analysis confirmed increased OAS1/3 protein levels in Pfn1 KO cells (**Figure S6**). Consistent with these findings, we also detected significantly greater abundance of IFNβ (an early response type I IFN) in the culture supernatant of Pfn1 KO 4T1 cells vs their control counterparts (**Figure 4C**, standard curve shown in **Figure S7**). Upstream Regulator Analyses (URA) of our transcriptome data identified STAT1 and interferon regulatory factors (IRFs) to be within the top 4 activated transcription factors (TFs) when Pfn1 expression is lost in both 4T1 and MDA-231 cells (**Figure 4D; Table S1** lists the top 15 predicted activated and repressed TFs in response to Pfn1 loss in each of the two breast cancer cell lines). By immunoblot analyses, we confirmed upregulation of S727-phosphorylated form of STAT1 (denoted as pSTAT1 hereon) in MDA-231 and 4T1 cells upon Pfn1 loss; in 4T1 cells, this was also accompanied by an increase in the total expression level of STAT1, as well as upregulation of IRF7 and its phosphorylated form (pIRF7), a STAT1-inducible gene and TBK1 target protein (**Figures 4E-H**). Overall, these data support activation of a functional Type I IFN response triggered by Pfn1 loss in breast cancer cells.

**Figure 4.**
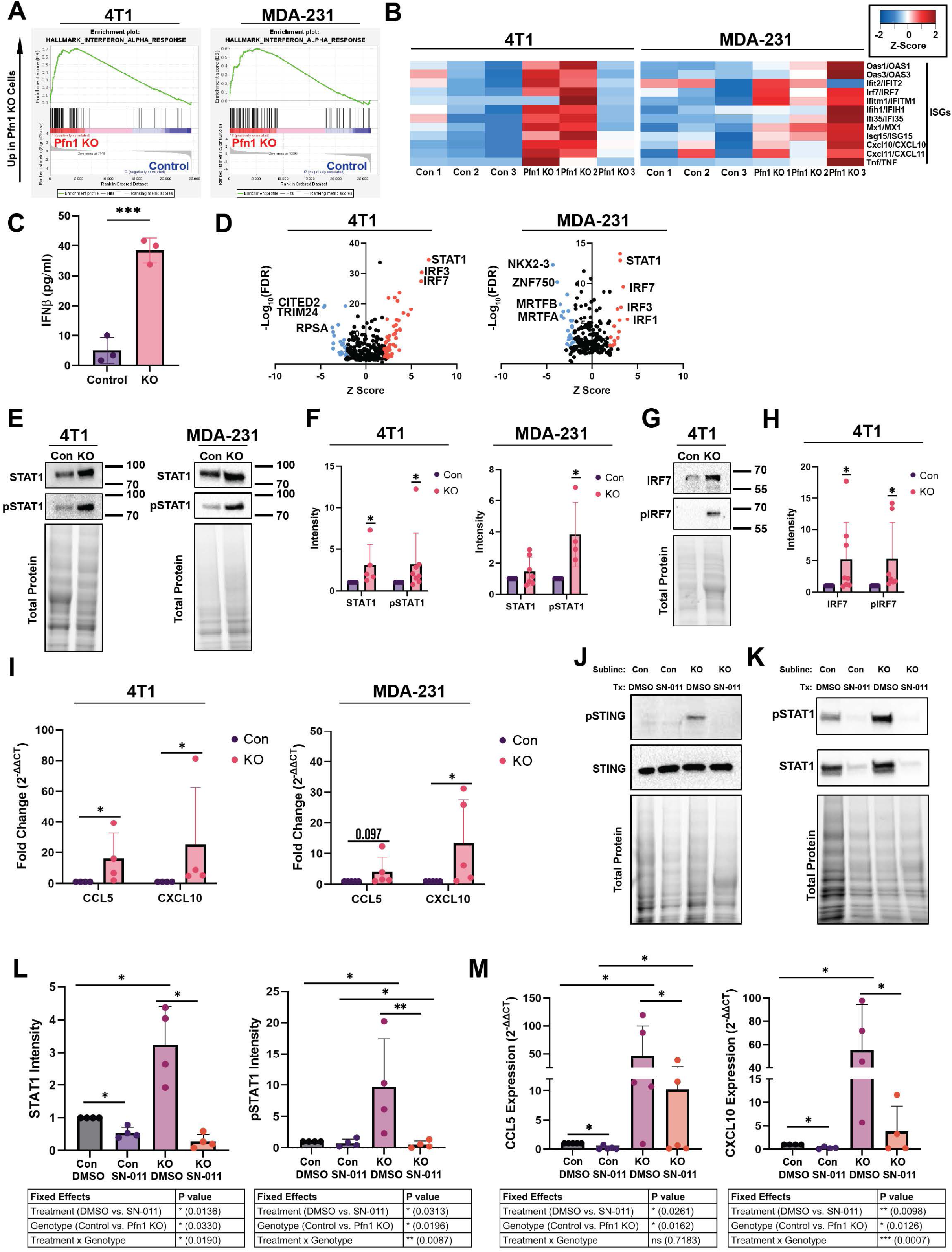
Pfn1 Loss Triggers a Type I Interferon Response. **A)** GSEA enrichment plots for the human and mouse HALLMARK_INTERFERON_ALPHA_RESPONSE gene set (4T1: nominal p value = <0.0001, normalized enrichment score = 1.502; MDA-231: nominal p value = 0.1933, normalized enrichment score = 1.407. **B)** Heat-plots showing Pfn1-dependent changes in the mRNA expression (represented as Z-scores) of major ISGs. **C)** Quantification of ELISA assay measuring IFNβ concentration in the supernatant of control and Pfn1 KO 4T1 cells. A student’s T test was used to compare means. **D)** Scatter plot showing IPA URA for transcription factors (TFs). TFs with the largest FC (positive or negative) are labeled on the plot. Cut off for significance (red/blue) was Z-score ≥ |2| and p-value < 0.05. **E, F)** Representative immunoblots and the associated quantification showing the relative expression levels of STAT1 and pSTAT1 (Ser727) in the indicated cell lines. A one-sample ratio T-test was used to compare experimental values to a hypothetical mean of 1. **G, H)** Representative immunoblots and the associated quantification showing the relative expression levels of IRF7 and pIRF7 (Ser 437/438) in the indicated cell lines. A one-sample ratio T-test was used to compare experimental values to a hypothetical mean of 1. **I)** qPCR results for CCL5 and CXCL10 in the indicated cell lines. A pair matched t test of the C_T_ values was used to compare between groups. **J)** Representative immunoblots for STING and pSTING (Ser365) demonstrating the inhibition of STING phosphorylation with SN-011 treatment. **K)** Representative immunoblot showing the expression levels STAT1 and pSTAT1 (Ser727) in control vs Pfn1 KO 4T1 cells with or without SN-011 treatment. **L)** Quantification of immunoblot analyses of STAT1 and pSTAT1 with or without SN-011 treatment in control and Pfn1 KO 4T1 cells. **M)** qPCR results for CCL5 and CXCL10 in control and Pfn1 KO 4T1 cells with or without SN-011 treatment in control and Pfn1 KO 4T1 cells. A mixed-effects REML model was used to evaluate the effects of Drug (DMSO vs. SN-011), Genotype (Con vs. KO), and their interaction on STAT1, pSTAT1, CCL5, and CXCL10 levels in 4T1 cells, with experimental replicate included as a matching factor across both variables to control for inter-experimental variability. Effect significance is indicated in the figures. Simple effects were probed using uncorrected Fisher’s LSD, comparing means within each drug condition and each genotype. The Pfn1 KO group in all experiments involving 4T1 cells represents clone C#2. **** p < 0.0001, *** p < 0.001, ** p < 0.01, * p < 0.05.

Type I IFN response plays a key role in promoting both innate and adaptive immunity, partly through enhanced production of pro-inflammatory cytokines and chemokines, including CCL5 and CXCL10 – two major chemoattractants for T cell recruitment. By qRT-PCR analyses, we found that both genes were transcriptionally upregulated in 4T1 cells; likewise, CXCL10 was upregulated and CCL5 showed a trend towards increased expression in MDA-231 cells (**Figure 4I**). Since STING activation is known to promote expression/activation of STAT1 (46) as well as expressions of CCL5 and CXCL10 (48), we next asked whether these features of Pfn1-deficient breast cancer cells are dependent on the action of STING pathway. To address this, we treated control and Pfn1 KO 4T1 cells with either 10 µM SN-011 (a small-molecule STING inhibitor that blocks STING activation by preventing its oligomerization and downstream signaling (49)) or DMSO as vehicle control. By immunoblot analyses, we confirmed dramatic downregulation of STING phosphorylation by SN-011 treatment (**Figure 4J**). SN-011-mediated STING inhibition dramatically reduced STAT1/pSTAT1 protein levels and CCL5/CXCL10 mRNA expression in both control and Pfn1 KO cells. Notably, a significant interaction effect between Pfn1 KO and STING inhibition was observed for STAT1, pSTAT1, and CXCL10, indicating that STING inhibition abolished the genotype-dependent differences in these targets. This interaction effect was not significant for CCL5, suggesting that CCL5 upregulation in 4T1 Pfn1 KO cells may involve additional STING-independent regulatory mechanisms (**Figure 4K-M**). These results indicate STING signaling is a key upstream driver of STAT1 activation and chemokine upregulation in this context.

### Pfn1 loss in tumor cells drives a T -cell-mediated anti-tumor response in breast cancer

Next, to determine whether triggering Pfn1 loss in cancer cells has any immunological impact in the tumor microenvironment, we performed *in vivo* studies. Specifically, we orthotopically implanted control and uninduced (i.e. without prior exposure to dox) Pfn1 KO 4T1 cells into the fourth inguinal mammary fat pad of fully immunocompetent BALB/c mice and allowed the tumors to establish for 7 days prior to triggering Pfn1 KO through administration of dox-containing chow. Longitudinal monitoring of tumor growth revealed striking regression of Pfn1 KO tumors beginning approximately ten days after dox induction (**Figure 5A**), resulting in a dramatically lower endpoint tumor burden relative to the control group (**Figure 5B**). Immunohistochemistry analyses of tumors showed significantly greater tumor infiltration of T cells (including effector CD8^+^ T cells) in Pfn1 KO vs control tumors after 14 days of dox administration (**Figure 5C-D).** These findings were orthogonally corroborated by flow cytometry-based analyses of relative percentages of CD3^+^, CD3^+^CD4^+^ and CD3^+^CD8^+^ cells in a subset of control vs Pfn1 KO tumors at a midpoint of the experiment, 7 days after dox administration (**Figures 5E, S8**). Since we observed that Pfn1 KO promotes senescence but does not induce overt cytotoxicity in 4T1 cells in culture (**Figure S5**), we reasoned that the observed regression of Pfn1 KO tumors could be mediated by a T cell-mediated immune response rather than a direct consequence of Pfn1 loss on tumor cell viability. To test this, we initially repeated these tumor model studies in immunodeficient BALB/c nude mice which lack functional T cells. While the endpoint Pfn1 KO tumors were still modestly smaller than the control tumors (this is consistent with the proliferation defect induced by Pfn1 loss), notably, triggering Pfn1 KO did not cause tumor regression in an immunodeficient setting (**Figure 6A-B**). Second, to specifically investigate whether tumor regression induced by Pfn1 loss in an intact immune setting is mediated by the action of CD8^+^ T cells, we performed 4T1 tumor model studies in immunocompetent Balb/c mice with or without antibody-mediated CD8+ T cell depletion (isotype control antibody treatment served as the control arm in these studies), as schematically depicted in **Figure 6C**. Validation of successful depletion of CD8^+^ T cells is shown in **Figure S9**. Further validating the immune-dependent regression model of Pfn1 KO tumors, CD8^+^ depletion was sufficient to prevent the Pfn1 KO-induced tumor regression and abrogated the difference in tumor weight between the control and Pfn1 KO tumors (**Figures 6D-E**). These *in vivo* data are consistent with a scenario that T cell-mediated tumor clearance is responsible for the robust regression phenotype of Pfn1 KO tumors observed in an intact immune setting. Collectively, our *in vitro* and *in vivo* findings support the notion that loss of Pfn1 leverages cGAS/STING signaling to trigger type I IFN response and T cell-mediated anti-tumor immune response in breast cancer (**Figure 7**).

**Figure 5.**
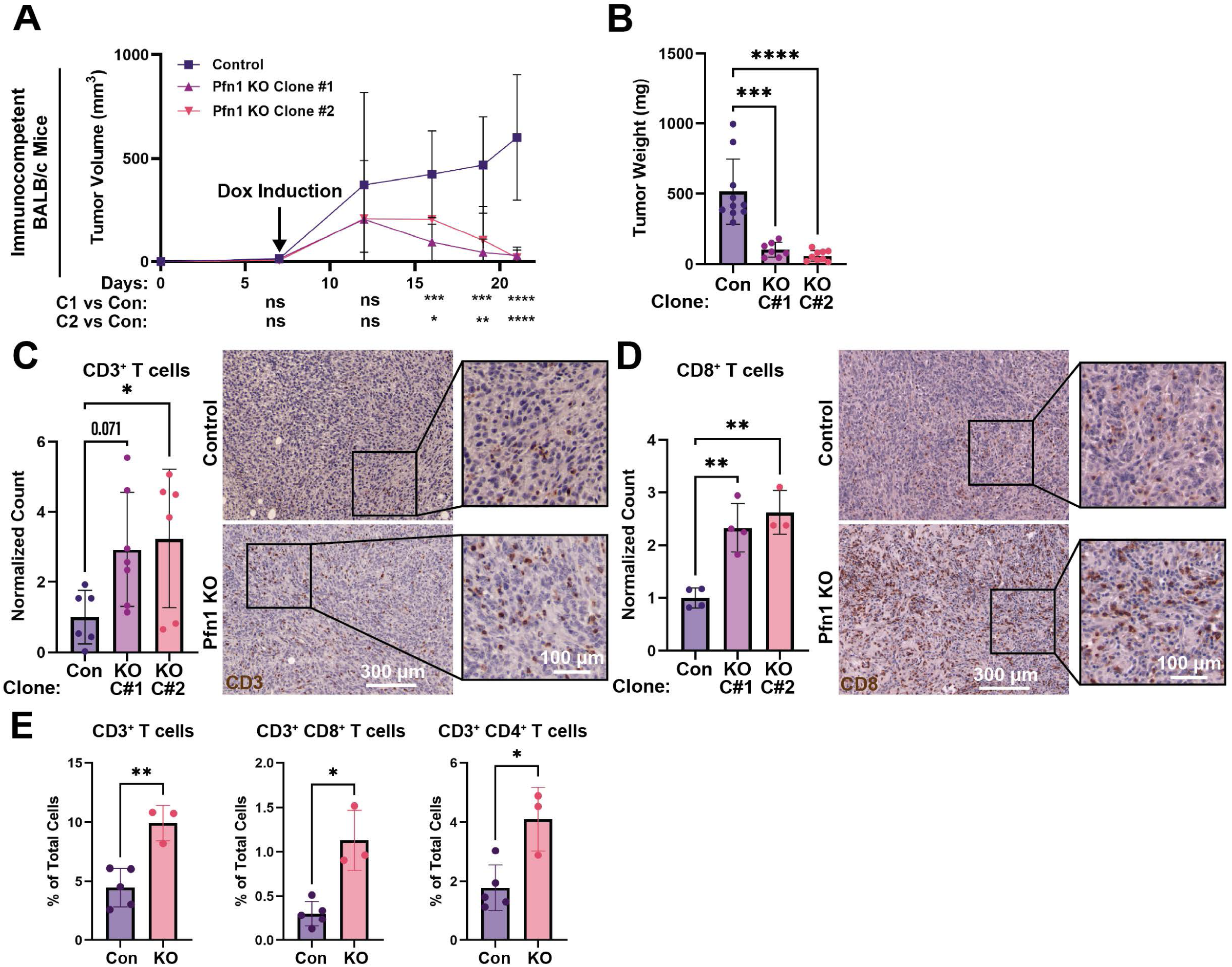
Pfn1 Deficiency Drives Tumor Regression and T Cell Recruitment. **A)** Growth kinetics of control and Pfn1 KO 4T1 orthotopic tumor grafts in fully immunocompetent Balb/c mice. Multiple t tests with Benjamini-Hoechberg correction and FDR < 0.01 were used to compare between groups at each time point. **B)** Summary of endpoint (21 days post-inoculation) 4T1 tumor burden representing the indicated groups. Data pooled from three independent experiments (number of mice: con = 10, Pfn1 KO (C#1) = 8, Pfn1 KO (C#2) = 8). One-way ANOVA with a post-hoc Tukey’s test was used to compare the average end-point tumor burden between the groups. **C)** Quantification of the number of CD3^+^ cells per 10X field and representative images of 10X fields of tumors (21 days post inoculation) DAB-stained for CD3. Number of tumors analyzed: Con = 6, Pfn1 KO (C#1) = 7, Pfn1 KO (C#2) = 6 with multiple (3–5) fields imaged per tumor. A one-way ANOVA with post-hoc Tukey’s test was used to compare between groups. **D)** Quantification of the number of CD8^+^ cells per 10X field and representative images of 10X fields of tumors (21 days post inoculation) DAB-stained for CD8. Number of tumors analyzed: Con = 4, Pfn1 KO (C#1) = 4, Pfn1 KO (C#2) = 3 with multiple (3–5) fields imaged per tumor. A one-way ANOVA with post-hoc Tukey’s Test was used to compare between groups. Pfn1 KO C#1 4T1 tumors are used as the representative image for Pfn1 KO tumors in both Figure 5C and 5D. **E)** Quantification of flow cytometry data of the percent of live singlet cells which are CD3^+^, CD3^+^CD4^+^, or CD3^+^CD8^+^ in tumors (14 days post inoculation). Con = 5 tumors, Pfn1 KO = 3 tumors. A student’s T-test was used to compare between groups. Pfn1 KO Clone #2 4T1 cells were used as the Pfn1 KO clone in this analysis. **** p < 0.0001, *** p < 0.001, ** p < 0.01, * p < 0.05.

**Figure 6.**
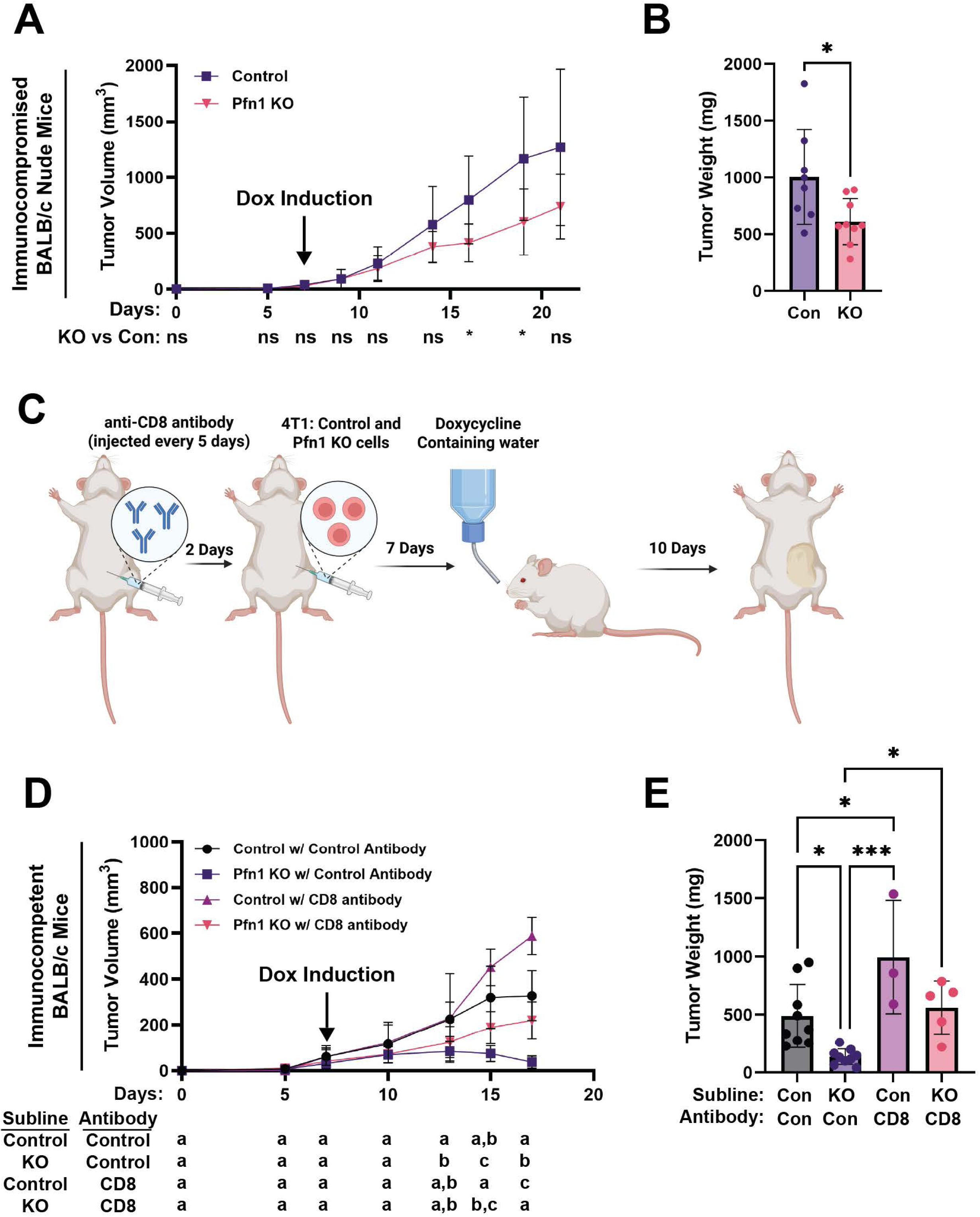
Pfn1 deficiency-induced tumor regression depends on an intact immune system. **A)** Growth kinetics of control and Pfn1 KO 4T1 (clone C#2) orthotopic tumor grafts in Nude Balb/c mice. Multiple t tests with Benjamini-Hoechberg correction and FDR < 0.01 were used to compare tumor burden between groups at each time point. **B)** Summary of endpoint (21 days post-inoculation) tumor burden for the indicated groups (data based on 8 control and 9 Pfn1 KO 4T1-inoculated mice pooled from two independent experiments). A student’s T test was used to compare the means. **C)** Schematic depicting the experimental procedure adopted for the CD8^+^ T cell depletion studies. **D)** Growth kinetics of control and Pfn1 KO (clone C#2) 4T1 tumors in immunocompetent Balb/c mice with or without CD8^+^ T cell depletion. Tumor volumes were analyzed using a two-way mixed ANOVA. A Tukey’s test was used to compare all four treatment groups at each time point. Compact letter display was used to report the results of these comparisons. Specifically, groups that share at least one letter are not significantly different from each other, while groups that share no letters are significantly different. **E)** Summary of endpoint (17 days post-inoculation) tumor burden for the indicated groups pooled from 2 experiments (n=9 (control cells, control antibody); n=3 (control cells, CD8 antibody); n=9 (Pfn1 KO cells, control antibody); n=5 (Pfn1 KO cells, CD8 antibody); the lower sample size in control and Pfn1 KO 4T1-bearing mice subjected to CD8+ T cell depletion resulted from early moribund status of these animals. One-way ANOVA with a post-hoc Tukey’s test was used to compare groups. **** p < 0.0001, *** p < 0.001, ** p < 0.01, * p < 0.05.

**Figure 7.**
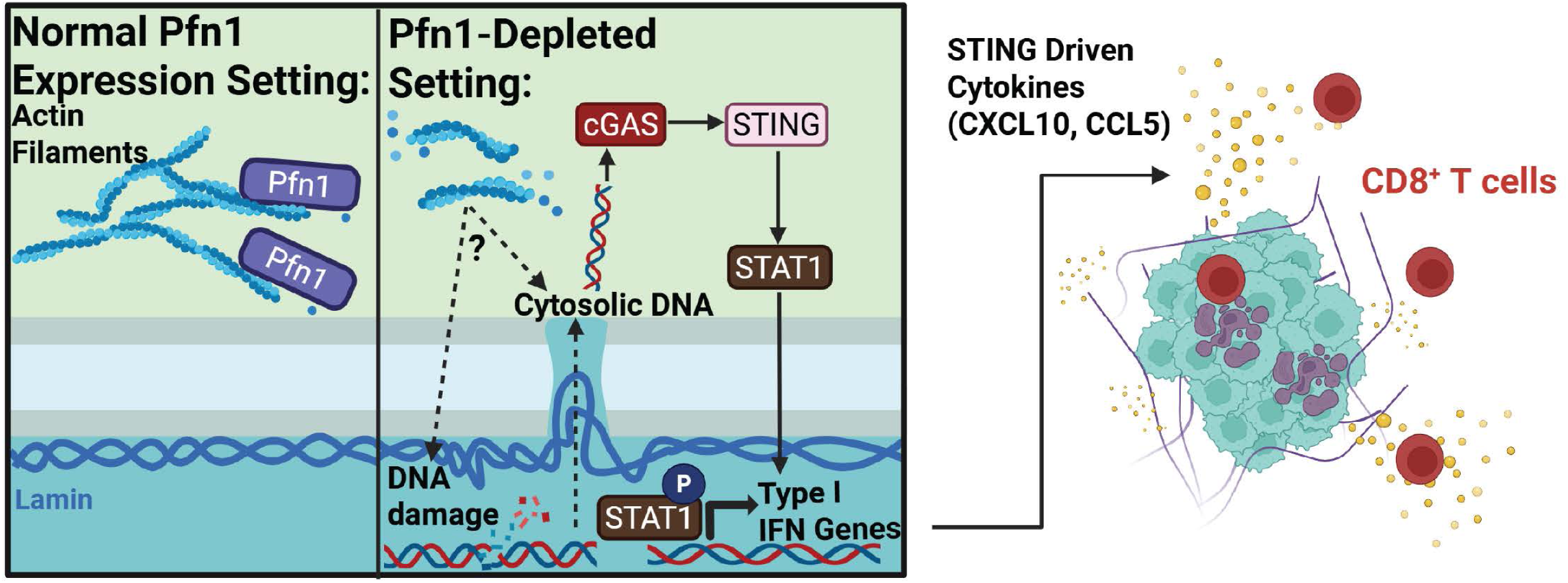
Schematic of the proposed mechanism of Pfn1-dependent regulation of inflammatory signaling. We propose that through a reduction in the polymerization of actin filaments (or disruption of Pfn1’s interaction with other non-actin proteins), the depletion of Pfn1 results in an increase in DNA damage and subsequent cytosolic DNA accumulation. Mitotic defects arising from Pfn1 depletion likely further contribute to this accumulation. The accumulated cytosolic DNA triggers the DNA-sensing-protein cGAS leading to a STAT1-mediated type-1 interferon response driving the expression of T cell recruiting chemotactic factors. In vivo, these chemotactic factors drive CD8^+^ T cell infiltration and CD8^+^ T cell-mediated tumor regression.

## DISCUSSION

While there is emerging evidence for Pfn1’s role in DNA repair and maintenance of genomic integrity(20, 24–26), the implication of nuclear abnormality in the absence of Pfn1 in disease contexts is still unclear. This study is the first report that functionally links Pfn1 loss to nuclear abnormality, type I IFN response, and T-cell mediated anti-tumor immunity utilizing breast cancer as a model system. Specifically, we report four novel findings. First, we provide the first direct evidence for reduced capacity for DNA DSB repair via both HR and NHEJ mechanisms in Pfn1-depleted conditions, findings that we were further able to mirror in human cancer through demonstration of the highest fraction of genomic alteration associated with the lowest quartile of Pfn1 expression. Second, we demonstrate that cellular Pfn1 loss potentiates the formation of rupture-prone micronuclei, cytosolic leakage of DNA, and cGAS-STING activation. Third, consistent with STING activation, we demonstrate that Pfn1 depletion in cancer cells promotes a type I IFN response and STING-dependent STAT1 activation, fostering a T cell chemoattractant milieu. Fourth, we demonstrate that triggering Pfn1 loss in cancer cells drives intratumoral infiltration by T cells and CD8^+^ T cell-mediated tumor regression in an orthotopic mouse model of breast cancer, further underscoring immunological benefit of Pfn1 depletion in cancer cells *in vivo*.

A previous study showed that Pfn1 is actively recruited from the cytoplasmic pool to the spindle midzone during anaphase. This recruitment appears to supply actin filaments to the forming cleavage furrow (20). Its loss therefore impairs the acto-myosin contractile ring needed for cytokinesis. As a result, Pfn1 loss leads to various mitotic errors including multipolar spindles, chromosome misalignment, anaphase bridges, lagging chromosomes, and cytokinesis defects resulting in binucleated/tetraploid daughter cells. Two additional studies also independently established that Pfn1 loss causes failed abscission and nuclear anomalies (multinucleated cells with irregular nuclei, abnormal mitotic spindles and mitotic catastrophe) in other cellular contexts (6, 25). These abnormalities likely explain the catastrophic consequence of Pfn1 loss during mammalian embryonic development where lethality ensues at the 2-cell stage (50). These findings resonate with our observation of the polyploidy phenotype of Pfn1 KO breast cancer cells. Both breast cancer cell lines in Pfn1-deficient condition appeared to be tolerant to multi-nucleation likely due to loss of functionality of the p53 gene in both cell lines (51, 52). Interestingly, Pfn1 overexpression also increases mitosis timing (20) and has a tumor-suppressive effect on breast cancer cells (22, 23), suggesting that a balanced level of Pfn1 is ideal for optimal cell cycle.

Beyond its canonical cytoplasmic actin-related functions, there is emerging evidence implying Pfn1’s role in DNA repair. For example, it has been reported that Pfn1-deficient cells when exposed to genotoxic agents exhibit persistent γH2AX accumulation and an impaired ability to resolve NHEJ-repair protein 53bp1 foci (26). While suggestive, these studies did not directly address whether the intrinsic DNA DSB repair capacity by either HR or NHEJ is affected by loss of Pfn1. By utilizing pathway-specific reporter systems, we herein demonstrate for the first time that loss of Pfn1 has a negative impact on both HR- and NHEJ-mediated DSB repair. This may explain why Pfn1-deficient cells even in the absence of any genotoxic stress are prone to accumulating DNA damage. Consistent with this, tumors with low Pfn1 expression exhibit the highest fraction of genome alteration in human cancer (as shown in this study), and Pfn1-haploinsufficiency has previously been associated with increased chromosome-level copy number alterations in osteosarcoma cells (20). Although the exact underlying mechanisms of how loss of Pfn1 leads to reduced efficiency of DNA repair are unclear, we can consider several possibilities. First, although Pfn1 is primarily a cytosolic protein, a small intracellular pool of Pfn1 also resides in the nucleus and serves as a key mediator of promoting nuclear export of actin through an interaction with exportin-6 (53). Given previous evidence for rapid nuclear translocation of Pfn1 in response to genotoxic stress and resolution of 53bp1-positive foci in Pfn1-deficient astrocytes upon re-expression of nuclear-restricted Pfn1 (26), one would presume that nuclear Pfn1 directly participates in DSB repair. By being a key regulator of nuclear actin and interactor of several important actin nucleation-promoting factors (NPFs – e.g. Arp2 and formin-family protein FMN2) that are important for DSB repair (54, 55), nuclear Pfn1 may facilitate DSB repair through modulating nuclear actin dynamics. Second, DNA repair can also be negatively impacted by replication stress, and Pfn1 has an emerging role in regulating replication fork dynamics. A recent study has identified that Pfn1 binds to SNF2H, a poly-proline domain containing nucleosome remodeler (56). Under normal conditions, Pfn1-SNF2H interaction increases DNA replication initiation and replication fork progression. However, under replication stress induced by hydroxyurea treatment (depletes dNTPs), elevating (specifically nuclear) Pfn1 level leads to more frequent replication fork stalling in part through inhibition of BOD1L (another polyproline domain-containing interactor of Pfn1) negatively affecting genome stability. While these overexpression-related findings are at apparent odds with greater accumulation of DNA damage in Pfn1-deficient condition reported in our and other studies, we also believe that a direct comparison is not appropriate here since Pfn1’s ability to promote replication fork stalling is applicable to specific experimental conditions (i.e. replication stress). In fact, since Pfn1’s action promotes replication fork progression under normal conditions, it is possible that loss of Pfn1-SNF2H interaction in Pfn1-depleted condition may exacerbate fork stalling and impair DNA repair process, accounting for an actin-independent DNA repair regulation by nuclear Pfn1. Third, consistent with Pfn1’s positive correlation with DNA repair pathways in clinical breast cancer samples reported by another group (57), we have found that Pfn1 loss impacts the expression of several major DNA repair-associated proteins. Pfn1 loss leads to coordinated downregulation of the chromatin remodeling factors SMARCAD1, KAT2, and SETMAR, which are required to relax nucleosomes flanking DSBs and make the underlying DNA accessible to repair enzymes. Transcriptional downregulation of end-resection nucleases (RBBP8, DNA2, and EXD2) and the HR mediator RAD52 could be additional contributing factors for reduced HR proficiency of Pfn1-deficient cells. Fourth, loss of Pfn1 may foster an intracellular environment that is inhibitory for DNA repair. Along this line, we have unpublished observations that cellular reactive oxygen species (ROS) level is prominently increased upon loss of Pfn1 expression. This observation parallels previous findings that Pfn1 KO increases the level of mitochondrial superoxide (10). Since ROS can induce DNA damage as well as impair the repair machinery by oxidative modification (58), Pfn1 loss can negatively impact DNA repair process by increased accumulation of ROS. These different possibilities should be explored in the future to advance our understanding of Pfn1’s role in DNA repair.

A key finding of our study is that Pfn1 loss promotes leakage of nuclear DNA and cGAS-STING activation. Micronuclei, particularly those without proper lamin enclosure, are vulnerable to rupture causing cGAS-STING activation. Although a previous study also reported increased presence of micronuclei in Pfn1-deficient osteoblasts(20), those micronuclei were not structurally characterized. Our studies show that Pfn1-deficient cells not only exhibit an increased frequency of micronuclei formation but also show a trend toward a greater proportion of those micronuclei completely lacking lamin-B1 in their envelopes, Therefore, micronuclei in Pfn1-depleted cells might be structurally unstable. However, this was not attributed to general downregulation of lamin abundance in the nuclear envelope. In fact, we found that Pfn1 KO cells display an increase in the overall contents of both lamin-B and -A/C at the nuclear lamina but at the same time exhibit features that are indicators of nuclear envelope instability (nuclear asymmetry and nuclear membrane invagination) mimicking phenotypes induced by overexpression of ALS-linked mutants of Pfn1 previously reported by another group (59). Because some of these ALS-linked Pfn1 mutations impair actin-related function, we speculate that the lamin abnormalities seen with Pfn1 knockout result from actin cytoskeletal dysfunction. We note, however, that these mutations also cause other significant changes in Pfn1 function, most notably an increased propensity to aggregate (60). Also, notably loss of Lamin-B1 is a common feature of cellular senescence (61), however in our study Pfn1 KO cells exhibited increased Lamin-B1 and increased SA-β-gal staining. However, it has been reported that under certain conditions (oxidative stress) senescence can also be associated with elevated Lamin-B1(62, 63).

Important downstream consequences of STING activation are type I IFN response, STAT1 activation, and production of T cell-supportive chemokines that are directly or indirectly regulated by STAT1 (CCL5 and CXCL10), all of which are recapitulated in Pfn1-deficient breast cancer cells in our studies. These *in vitro* findings are also consistent with T cell-mediated anti-tumor immune response triggered by Pfn1 loss in breast cancer cells *in vivo*. Disrupted actin cytoskeleton is a hallmark feature of Pfn1-deficient cells. One could consider a complex interplay between Pfn1, STING, and STAT1 signaling involving modulation of actin cytoskeleton. Since F-actin collapse can lead to nuclear deformation, cytoplasmic DNA accumulation, and cGAS/STING activation (64), one likely explanation is that through disruption of actin cytoskeleton, Pfn1 loss promotes cGAS/STING activation leading to IRF7-driven upregulation of STAT1 expression as well as its activation (via induction of type 1 IFNs). Interestingly, there is also evidence for crosstalk between STING pathway and actin remodeling, and actin-dependent modulation of STAT1 activation. For example, it has been shown that STING deficiency can promote abnormal F-actin accumulation in B cells in a WASP-dependent manner (65). Other studies reported that actin disruption by pharmacological strategies prolongs STAT1 activation (but not the expression), mainly by regulating the conformation of tyrosine phosphorylated STAT1 that restricts its access to phosphatases (66, 67). Therefore, there might be additional contribution of a direct actin-dependent regulation of STAT1 functionality without STING’s involvement in Pfn1-depleted cells. Since our studies show that STING inhibition nearly abrogates Pfn1-dependent changes in STAT1 expression/activation and chemokine expression, we believe that STING activation is the dominant mechanism of for STAT1- and chemokine-related effects elicited by Pfn1 loss in breast cancer cells. Future studies involving F-actin modulation in experimental settings with or without functional STING will shed further mechanistic insights.

The findings reported in this study have therapeutic implications in the context of cancer. First, genomic instability is a defining hallmark of breast cancer and a key driver of tumor evolution, therapeutic resistance, and immune recognition. Accumulation of DNA damage, particularly DNA DSBs, fuels chromosomal rearrangements, copy number alterations, and aneuploidy that shape tumor heterogeneity and clinical behavior. In triple-negative breast cancer (TNBC), high levels of chromosomal instability are frequently observed and are associated with aggressive disease and poor outcomes (68, 69). At the same time, genomic instability creates therapeutic vulnerabilities, as illustrated by the clinical success of PARP inhibitors in tumors harboring HR deficiencies (70). Thus, understanding how breast cancer cells tolerate or respond to DNA damage is central to both tumor biology and translational intervention. Our findings that Pfn1 loss leads to HR deficiency and increased DNA damage while low Pfn1 expression correlates with higher proportions of fraction of the genome altered in human cancer, indicates that Pfn1- or Pfn1-associated pathway targeting may be a potential method for enhancing PARP inhibition success or Pfn1 reduction in tumors may serve as a biomarker for PARP inhibition treatment evaluation. Second, although our *in vivo* findings are limited to one breast cancer cell line (a limitation of this study), it supports a premise that Pfn1 loss enhances tumor immunogenicity in breast cancer. If these results are reproducible in broader model settings of breast cancer, it could pave the way for further explorations of whether Pfn1 expression level in tumors serves as a biomarker for immunotherapy responsiveness, and/or targeting Pfn1 is an effective strategy to augment the therapeutic efficacy of immune checkpoint inhibitors in breast cancer. In general, identifying the functions of Pfn1 which contribute to STING activation and HR deficiencies would strategize targeting of Pfn1 or its associated effectors in these pathways.

In conclusion, this study establishes Pfn1 loss in tumor cells as a novel conceptual strategy of boosting STING activation and anti-tumor immunity in breast cancer. Many important questions remain unresolved which need to be addressed in future studies.

## MATERIALS AND METHODS

### Cell Culture and Genetic Manipulation

Doxycycline (Dox)-inducible Cas9-mediated Pfn1 knockout (KO) cells were generated from sublines of MDA-MB-231 and 4T1 cells that co-expressed GFP and Luciferase (gifts from Dr. Jennifer Koblinski, Virginia Commonwealth University) or from wild type 4T1 cells (for GFP- sublines). Dox-inducible Cas9 expression and constitutive Tet-on 3G expression were introduced via lentiviral transduction (Dharmacon; VCAS11227). These cells were subsequently transduced with lentivirus encoding one of two stably expressed Pfn1-targeting guide RNAs (gRNAs) - either Dharmacon VSGH10142-246635145 (Target Sequence: TGTCAGGACGCGGCATCGT; used for both MDA-231 and 4T1 cells) or Dharmacon VSGM10144-246796570 (Target Sequence: GATCTTCGTACCAAGAGCAC; used for 4T1 cells only). Non-targeting control gRNA (Dharmacon; VSGC12243) transduced cells served as control. For 4T1 cells, two clones of the GFP/Luciferase+ inducible Pfn1 KO cells were isolated and one clone of GFP/Luciferase- inducible Pfn1 KO cells was isolated. For 231 cells, one clone of inducible Pfn1 KO cells was isolated. MDA-231, 4T1 and HEK-293 cells were maintained and selected in DMEM (MDA-231, 4T1) or DMEM-F12 (HEK-293) supplemented with 10% fetal bovine serum, antibiotics/antifungals (100 U/ml penicillin, 100 µg/ml streptomycin, and 0.25 µg/ml Amphotericin B), and various selection markers including puromycin (2.5 µg/ml), blasticidin (10 µg/ml), and G418 (100 µg/ml). Selection markers were used during routine cell culture only and were omitted during experiments. To induce Pfn1 KO, cells were treated with 1 µg/ml Dox for 5 days prior to experimental evaluation, unless otherwise specified. To activate STING, 5 µg/ml 2’,3’-cGAMP (InvivoGen; Catalog #tlrl-nacga23-02) was transfected into cells using Lipofectamine 2000 (ThermoFisher; Catalog #11668019) overnight or cells were treated overnight with 1µM diABZI (Invivogen Catalog# tlrl-diabzi-2), a STING activator. For STING inhibition studies, STING inhibitor SN-011 (MedChemExpress; Catalog #2249435-90-1) was added directly to the media at 10 µM for 5 days, concurrent with Pfn1 KO induction.

For transient Pfn1 knockdown studies, HEK293 cells were seeded at 1 × 10⁵ cells/well in the wells of a 6-well plate. Wells were pre-coated with 10 µg/mL type I collagen (Gibco, A10483-01; 4 mg/mL stock diluted in ice-cold serum-free medium) and washed 2X with PBS prior to cell seeding. Twenty-four hours after seeding, cells were transfected with either 10 nM Pfn1-targeting siRNA (Dharmacon, custom siRNA strand 5’-AGAAGGUGUCCACGGUGGUUU-3’) or equivalent amount of non-targeting control siRNA (Dharmacon, D-001206-13-05) using Lipofectamine RNAiMAX (Thermo Fisher Scientific, #13778075) according to the manufacturer’s instructions as previously described (71).

### Protein Extraction and Immunoblotting

Total cell lysate was extracted using Pierce IP Lysis Buffer (Thermo Fisher Scientific; Catalog #87787) supplemented with 0.1% sodium dodecyl sulfate (SDS) and protease/phosphatase inhibitor cocktail (Thermo Fisher Scientific, Catalog #78440) and denatured by boiling in 1X Laemmli buffer prior to running SDS-PAGE. The details of all antibodies used for immunoblotting are listed in the *Supplementary Information* (**Table S2**). Total Protein loading was estimated for the purpose of band intensity normalization by imaging gels made with TGX Stain-Free FastCast Acrylamide Solution (BioRad, Catalog #1610180) after 1 minute UV activation.

### Immunostaining

Cells cultured in tissue culture-treated wells were washed with PBS, fixed in 3.7% paraformaldehyde (5 minutes), permeabilized with 0.5% Triton-X (10 minutes), and blocked with 10% goat serum (1 hour). Following overnight incubation at 4°C with the various primary antibodies (details are listed in *Supplementary Information* **Table S2**), cells were washed 3X with PBS, incubated with the appropriate fluorescent secondary antibodies (1:100, 1 hour, room temperature), washed, and then stained with DAPI and/or Phalloidin (Cytoskeleton Inc.; Catalog #PHDN1). Fluorescence image acquisition was performed on an Olympus IX83 inverted microscope with Cicero confocal attachment platform. Images were processed with Fiji. CellProfiler was used to develop custom analysis pipelines for measuring the Lamins intensity in the lamina and counting the number and intensity of γH2AX/RAD51/53BP1 foci.

### RNA-sequencing and Analysis

For bulk RNA sequencing, control and Pfn1 KO sublines of breast cancer cells were treated with doxycycline for 5 days prior to RNA collection with the Qiagen RNeasy Mini Plus Kit (Qiagen, Catalog # 74134) following the manufacturer’s protocol. The methodological details of RNAseq are described elsewhere (9). Using the transcript per million normalized gene expression values, GSEA enrichment plots were generated using the Broad Institute GSEA 4.3.3 software. Heatplots of the interferon stimulated genes (ISGs) were plotted using z-scores calculated by: (x – µ)/σ, where x is the value of a given gene for a given sample, µ is the mean value of a given gene across all samples, and σ is the standard deviation of a given gene across all samples. Upstream Regulator Analysis (URA) was performed using the Qiagen Ingenuity Pathway Analysis (IPA) platform.

### ELISA

To assess the IFN-β in the secretome of 4T1 cells ELISA was performed. Briefly, 4 days post doxycycline treatment of the cells, the culture media were changed to serum free media and cells were incubated for a further 24hrs. After this incubation period, the culture supernatant was collected from the cells and filtered with 0.45μM filter to remove any cell debris. The IFN-β ELISA were performed as per manufacture’s protocol (Abcam, Catalog #ab252363). The IFN-β concentration was calculated as per the standard curve.

### Animal Studies

All animal studies were conducted in compliance with an approved IACUC protocol and University of Pittsburgh Division of Laboratory Animal Resources guidelines. For orthotopic tumor graft studies, 1 × 10⁵ 4T1 cells (control or Pfn1 KO) were suspended in 50 µl of 1:1 PBS/Matrigel and inoculated into the fourth inguinal mammary fat pad of anesthetized 5–6-week-old female syngeneic BALB/c or BALB/c Nude (CAnN.Cg-Foxn1nu/Crl) mice (Jackson Labs). Seven days post-inoculation, mice were fed doxycycline-containing chow (Fisher Scientific; Catalog #14-726-309) for 5 days to induce Cas9-mediated Pfn1 KO. For CD8^+^ T cell depletion studies, mice were injected with 200µg of CD8 antibody (BioXCell, Catalog #BE0004-1) or Control IgG (BioXCell, Catalog #BE0090) on days -2, 3, 8, and 13 relative to tumor inoculation as done previously (72). Tumor volume was monitored by caliper measurements throughout and estimated using the formula V = 0.5 × L × W², where L and W are the longest and shortest dimensions, respectively. Mice were euthanized 14, 17, or 21 days post-inoculation (as specified in individual experiments) and tumors were harvested. For histology, tumors were fixed overnight in 4% paraformaldehyde, transferred to 70% ethanol overnight, and then paraffin-embedded and sectioned by the University of Pittsburgh Histology Core. Immunohistochemistry was performed using a rabbit-specific HRP/DAB detection kit (Abcam; Catalog #ab64264) with antibodies specified in the *Supplementary Information* (**Table S2**). For flow cytometry, tumors were manually homogenized and enzymatically digested with 100 U/ml collagenase at 37°C. Cells were stained with antibodies specified in the *Supplementary Information* (**Table S2**), followed by dead cell exclusion with propidium iodide (1 µg/ml) and fixation in 1% paraformaldehyde. Flow cytometry was performed on an Attune NxT Acoustic Focusing Cytometer and data were analyzed using FlowJo v11.

### Quantitative RT-PCR

Total RNA was extracted for cells in a six-well plates using the RNeasy mini kit (Qiagen, catalog # 74104) according to manufacturer’s protocol. Complementary DNA (cDNA) was synthesized from 1 µg of RNA using the Quantitect Reverse Transcription Kit (Qiagen, catalog # 205311) in total reaction volume of 20µl. Each PCR was prepared with 1µl cDNA, 1µM (CCL5/CXCL10) or 10nM (18S) forward and reverse primers, 12.5 µl of SYBR Select Master Mix (Thermo Fisher, catalog # 4472903) and water for a total reaction volume of 25µl. Quantitative RNA expression was measured by real-time PCR using the StepOnePlus system and StepOne software (Applied Biosystems). PCR cycling conditions were as follows: 95°C for 30 seconds, 50°C for 30 seconds, and 72°C for 1 minute, for 45 cycles. Primer sequences are provided in *Supplementary Information* (**Table S3**).

### DNA Repair Analysis

HR and NHEJ repair efficiencies were quantitatively assessed using the pDR-GFP reporter (Addgene #26475) (35) and the pLCN-DSB reporter (Addgene #98895) (73) assays, respectively, with DSBs induced by the I-SceI expression plasmid pCBASceI (Addgene #26477) (74). Specifically, 1 × 10⁵ HEK-293 cells (control vs Pfn1 knockdown) and 1.2 × 10⁵ 4T1 cells (control vs Pfn1 KO) were seeded in the wells of 6-well plate forty-eight hours after siRNA transfection and five days after doxycycline induction, respectively. Twenty-four hours later, cells were transfected with either a) pDR-GFP (2 µg) plus pCBASceI (2 µg) or b) pLCN-DSB (2 µg) plus pCBASceI (2 µg) or c) EGFP (Clontech-BD Biosciences, 2 µg) using Lipofectamine 2000 (Thermo Fisher Scientific, #11668027) according to the manufacturer’s instructions. Untransfected cells served as control. Forty-eight hours post-transfection, cells were harvested, fixed in 4% paraformaldehyde in PBS for 15 min at room temperature, washed, and resuspended in PBS for flow cytometry analysis on an Attune NxT Acoustic Focusing Cytometer, with at least 15,000 single-cell events recorded per sample. Viable singlets were selected by sequential FSC-H/SSC-H and SSC-A/SSC-H gating. GFP⁺ gates were defined using negative control cells (to establish the background) and EGFP-transfected cells (marked the positive population) with data analyses performed with the FlowJo (v10) software. For assessing repair efficiency, percentages of GFP⁺ cells in the reporter assays were normalized by matched % counts of EGFP-expressing cells (to correct for transfection efficiency of the reporter), and the values estimated for Pfn1 KO/knockdown groups were then normalized to the same estimated for the corresponding control groups for comparison.

### Cytosolic DNA Analysis

Cytosolic DNA was quantified following a published protocol (75). Briefly, approximately 4 million cells were trypsinized and resuspended in lysis buffer (150 mM NaCl, 50 mM HEPES, 25 µg/ml Saponin). After reserving one-third of this lysate as the total cell fraction, the remaining two thirds were processed to obtain the cytosolic fraction by sequential centrifugation: first at 1,000 × g to pellet intact nuclei and mitochondria, then at 17,000 × g to remove residual cell debris, retaining the clarified supernatant each time. Cytosolic DNA was extracted from the cytosolic fraction using the QIAquick Nucleotide Removal Kit (Qiagen, Catalog #28306) per the manufacturer’s instructions. Total cellular DNA was extracted from the whole cell fraction using the QIAamp DNA Micro Kit (Qiagen, Catalog #56304) per the manufacturer’s instructions. Quantitative PCR reactions were prepared using both DNA fractions. Each 25 µl reaction contained 1 µl of isolated DNA, 12.5 µl of SYBR Select Master Mix (Thermo Fisher, Catalog #4472903), 10 pmol/µl each of forward and reverse primers (See **Table S3** for primer information), and nuclease-free water to volume. Thermal cycling was performed under the following conditions: 95°C for 15 s and 60°C for 60 s, for 40 cycles. The relative abundance of cytosolic DNA was calculated using a modified 2^ΔΔCT^ method. ΔC_T_ was first determined by subtracting the average C_T_ of the cytosolic fraction from the average C_T_ of the whole cell fraction. ΔΔC_T_ was then calculated by subtracting the ΔC_T_ of the control group from the ΔC_T_ of the experimental group, and the result was used as the exponent (2^ΔΔC^_T_) to obtain the fold difference.

### Clinical Data Analysis

From the TCGA Pan-Cancer Atlas, mRNA expression (z-scores relative to normal samples) and corresponding clinical data, including fraction genome altered (FGA), age, sex, and cancer type, were obtained from the cBioportal portal. The cohort was composed of the following tumor types: Adrenocortical Carcinoma (n = 92), Cholangiocarcinoma (n = 36), Bladder Urothelial Carcinoma (n = 411), Colorectal Adenocarcinoma (n = 594), Breast Invasive Carcinoma (n = 1,084), Brain Lower Grade Glioma (n = 514), Glioblastoma Multiforme (n = 592), Cervical Squamous Cell Carcinoma (n = 297), Esophageal Adenocarcinoma (n = 182), Stomach Adenocarcinoma (n = 440), Uveal Melanoma (n = 80), Head and Neck Squamous Cell Carcinoma (n = 523), Kidney Renal Clear Cell Carcinoma (n = 512), Kidney Chromophobe (n = 65), Kidney Renal Papillary Cell Carcinoma (n = 283), Liver Hepatocellular Carcinoma (n = 372), Lung Adenocarcinoma (n = 566), Lung Squamous Cell Carcinoma (n = 487), Diffuse Large B-Cell Lymphoma (n = 48), Acute Myeloid Leukemia (n = 200), Ovarian Serous Cystadenocarcinoma (n = 585), Pancreatic Adenocarcinoma (n = 184), Mesothelioma (n = 87), Prostate Adenocarcinoma (n = 494), Skin Cutaneous Melanoma (n = 448), Pheochromocytoma and Paraganglioma (n = 178), Sarcoma (n = 255), Testicular Germ Cell Tumors (n = 149), Thymoma (n = 123), Thyroid Carcinoma (n=500), Uterine Corpus Endometrial Carcinoma (n = 529), and Uterine Carcinosarcoma (n = 57). Tumor purity estimates were derived from immunohistochemistry-based measurements as described previously (76). Age, sex, tumor purity, and cancer type were controlled for using a multivariable linear regression framework prior to association testing. Specifically, fractional genome alteration (FGA) values were modeled with these variables as covariates, and PFN1-associated effects were evaluated using the residualized values to remove variance attributable to demographic factors, tumor purity, and cancer-type–specific differences. Samples with missing values were excluded. The importance of correcting for purity when correlating gene expression and markers of genomic instability has been detailed previously (77, 78).

### Statistics

All statistical tests were performed in Graphpad Prism Software. Specifics of statistical tests are indicated in the figure legends. All data are presented as mean ± SD. For qPCR, all statistical analyses were performed on raw ΔCt values using paired T tests. For graphical representation, data are displayed as relative expression using the 2^−ΔΔCT^ method, normalized to the relevant control group. For immunoblots, all statistical analysis were performed on band intensity normalized to the total protein loading and then normalized to the relevant control for each given experiment. For comparisons between two groups in which the control was normalized to 1 (i.e. western blots, cytosolic DNA leakage experiments), one-sample t-tests were used to determine whether the experimental group differed significantly from the hypothetical mean of 1. For experiments involving two independent variables, a two-way ANOVA was performed using a mixed-effects analysis to account for missing data points where applicable. Post-hoc multiple comparisons were corrected using Tukey’s.

## AUTHOR CONTRIBUTIONS

IE conceptualized studies, performed experiments, analyzed data, wrote the manuscript and acquired funding; MB: performed experiments, analyzed data, and wrote manuscript; SKM: performed experiments, analyzed data, and wrote manuscript, VY, EW, AK: performed experiments and analyzed data; JL: analyzed data, SL, JL: oversaw experiments and data analyses; WJS – provided intellectual input and assisted with conceptualization of studies; PR: wrote/edited the manuscript, acquired funding, and was responsible for overall supervision of the project.

## CONFLICT OF INTEREST

The authors declare no conflict of interest.

## DATA AVAILABILITY STATEMENT

All primary data are included in the main manuscript and supplementary information. The transcriptomic data generated as a part of this study will be uploaded to the NCBI Sequence Read Archive (SRA). Any code used for image analysis or clinical data analysis will be available upon request

## ACKNOWLEDGEMENTS

Ian Eder was supported by NCI F31-CA306130, NIBIB T32-EB001026, NCATS TL1-TR001858. Work in the Roy lab was supported by grants from NIH (R01CA248873, R01CA271095, R21EY032632) and the Department of Defense (HT9425-24-1-0556).

---

**Table S1.** Top differentially activated transcription factors as predicted by IPA upstream regulator analysis.

| 4T1 Upregulated<br>(Pfn1 KO vs. Con) |  |  | 4T1 Downregulated<br>(Pfn1 KO vs. Con) |  |  | MDA-231 Upregulated<br>(Pfn1 KO vs. Con) |  |  | MDA-231 Downregulated<br>(Pfn1 KO vs. Con) |  |  |
| --- | --- | --- | --- | --- | --- | --- | --- | --- | --- | --- | --- |
| TF | Z-Score | -log<sub>10</sub><br>(FDR) | TF | Z-Score | -log<sub>10</sub><br>(FDR) | TF | Z-Score | -log<sub>10</sub><br>(FDR) | TF | Z-Score | -log<sub>10</sub><br>(FDR) |
| STAT1 | 6.94 | 34.50 | TRIM24 | -4.66 | 18.77 | IRF1 | 3.90 | 5.593 | NKX2-3 | -4.31 | 12.31 |
| IRF3 | 6.12 | 30.40 | CITED2 | -4.56 | 19.34 | IRF7 | 3.41 | 9.602 | ZNF750 | -3.8 | 10.18 |
| IRF7 | 6.07 | 27.42 | RPSA | -3.70 | 12.04 | IRF3 | 3.17 | 7.051 | MRTFB | -3.49 | 7.446 |
| ZBTB10 | 4.97 | 18.12 | IRF2BP2 | -3.60 | 5.226 | STAT1 | 3.08 | 12.86 | MRTFA | -3.37 | 6.021 |
| RELA | 4.67 | 16.27 | ETV3 | -3.60 | 10.59 | NONO | 3.06 | 13.69 | ETV3 | -3.16 | 6.537 |
| NONO | 3.76 | 11.20 | PRDM1 | -3.58 | 6.899 | PRDM2 | 3.00 | 5.463 | ETV6 | -3.11 | 5.723 |
| BHLHE40 | 3.71 | 11.45 | NKX2-3 | -3.52 | 9.304 | IRF5 | 2.94 | 4.057 | GATA4 | -2.73 | 2.455 |
| IRF1 | 3.68 | 23.65 | ETV6 | -3.50 | 9.376 | RUNX3 | 2.82 | 3.324 | RPSA | -2.72 | 8.669 |
| FOXO1 | 3.66 | 4.723 | MYB | -3.05 | 12.96 | IRF9 | 2.77 | 6.838 | TEAD2 | -2.64 | 4.177 |
| STAT4 | 3.59 | 14.46 | ZFP36 | -2.93 | 3.954 | SOX3 | 2.44 | 2.954 | MYC | -2.60 | 6.602 |
| NFATC2 | 3.47 | 20.92 | BCOR | -2.82 | 3.217 | GMNN | 2.44 | 2.860 | MEF2C | -2.57 | 3.018 |
| CEBPB | 3.37 | 14.66 | IRF4 | -2.81 | 7.414 | SOX1 | 2.44 | 2.436 | WBP2 | -2.53 | 3.348 |
| IRF5 | 3.35 | 11.05 | PPARGC1A | -2.74 | 3.924 | HDAC4 | 2.42 | 4.728 | TEAD3 | -2.44 | 3.191 |
| CTNNB1 | 3.31 | 12.96 | ETV5 | -2.71 | 4.124 | AIP | 2.23 | 2.252 | ATF4 | -2.42 | 2.202 |
| HIF1A | 3.32 | 8.573 | ETV7 | -2.64 | 6.438 | MYCN | 2.06 | 3.388 | TBX5 | -2.41 | 3.625 |

**Table S2.** Antibody Information.

| Target | Target Species | Clonality | Source | Catalog # | Concentration | Application |
| --- | --- | --- | --- | --- | --- | --- |
| phospho-STING (Ser365) | Mouse | monoclonal (Clone: D8F4W) | Cell Signaling | 72971 | 1:500 | Western Blotting |
| phospho-STING (Ser366) | Human | monoclonal (Clone: D7C3S) | Cell Signaling | 19781 | 1:250 | Western Blotting |
| STING | Mouse/Human | monoclonal (Clone: 9V9A3) | Invitrogen | MA5-51318 | 1:1000 | Western Blotting |
| STAT1 | Mouse/Human | Polyclonal | Cell Signaling | 9172 | 1:1000 | Western Blotting |
| phospho-STAT1 (Ser727) | Mouse/Human | monoclonal (Clone: D3B7) | Cell Signaling | 8826 | 1:1000 | Western Blotting |
| TREX1 | Mouse | monoclonal (Clone: EPR14985) | Abcam | ab185228 | 1:1000 | Western Blotting |
| IRF7 | Mouse | monoclonal (Clone: D8V1J) | Cell Signaling | 72073 | 1:500 | Western Blotting |
| phospho-IRF7 (S437/438) | Mouse | monoclonal (Clone: D6M2I) | Cell Signaling | 24129 | 1:500 | Western Blotting |
| GAPDH | Mouse | monoclonal (Clone: GA1R) | Invitrogen | MA5-15738 | 1:2000 | Western Blotting |
| Tubulin | Mouse | monoclonal (Clone: DM1A) | Invitrogen | 14-4502-82 | 1:2000 | Western Blotting |
| Profilin-1 | Mouse/Human | monoclonal (Clone: EPR6304) | Abcam | AB12904 | 1:4000 | Western Blotting |
| OAS1/3 | Mouse | Polyclonal | Proteintech | 14955-1-AP | 1:1000 | Western Blotting |
| IRF3 | Mouse | Monoclonal (Clone: D83B9) | Cell Signaling | 4302 | 1:100 | Immunofluorescence |
| γH2AX | Mouse | Polyclonal | ABClonal | AP0099 | 1:50 | Immunofluorescence |
| Lamin-B1 | Mouse | Polyclonal | Proteintech | 66095-1-Ig | 1:100 | Immunofluorescence |
| Lamin-A/C | Mouse | Monoclonal<br>(Clone: mab636) | Invitrogen | MA3-1000 | 1:100 | Immunofluorescence |
| 53BP1 | Mouse | Polyclonal | Invitrogen | PA5-27195 | 1:100 | Immunofluorescence |
| RAD51 | Mouse | Polyclonal | Invitrogen | PA1-16565 | 1:100 | Immunofluorescence |
| CD3 | Mouse | Monoclonal<br>(Clone: E4T1B) | Cell Signaling | 78588 | 1:100 | Immunohistochemistry |
| CD8 | Mouse | Monoclonal<br>(Clone: D4W2Z) | Cell Signaling | 98941 | 1:100 | Immunohistochemistry |
| APC CD3 | Mouse | Monoclonal<br>(17A2) | Invitrogen | 17-0032-82 | 1:50 | Flow cytometry |
| PerCP CD4 | Mouse | Monoclonal<br>(GK1.5) | Proteintech | CP-65104 | 1:50 | Flow cytometry |
| FITC CD8 | Mouse | Monoclonal (53-6.7) | Invitrogen | 11-0081-85 | 1:10 | Flow cytometry |

**Table S2.**
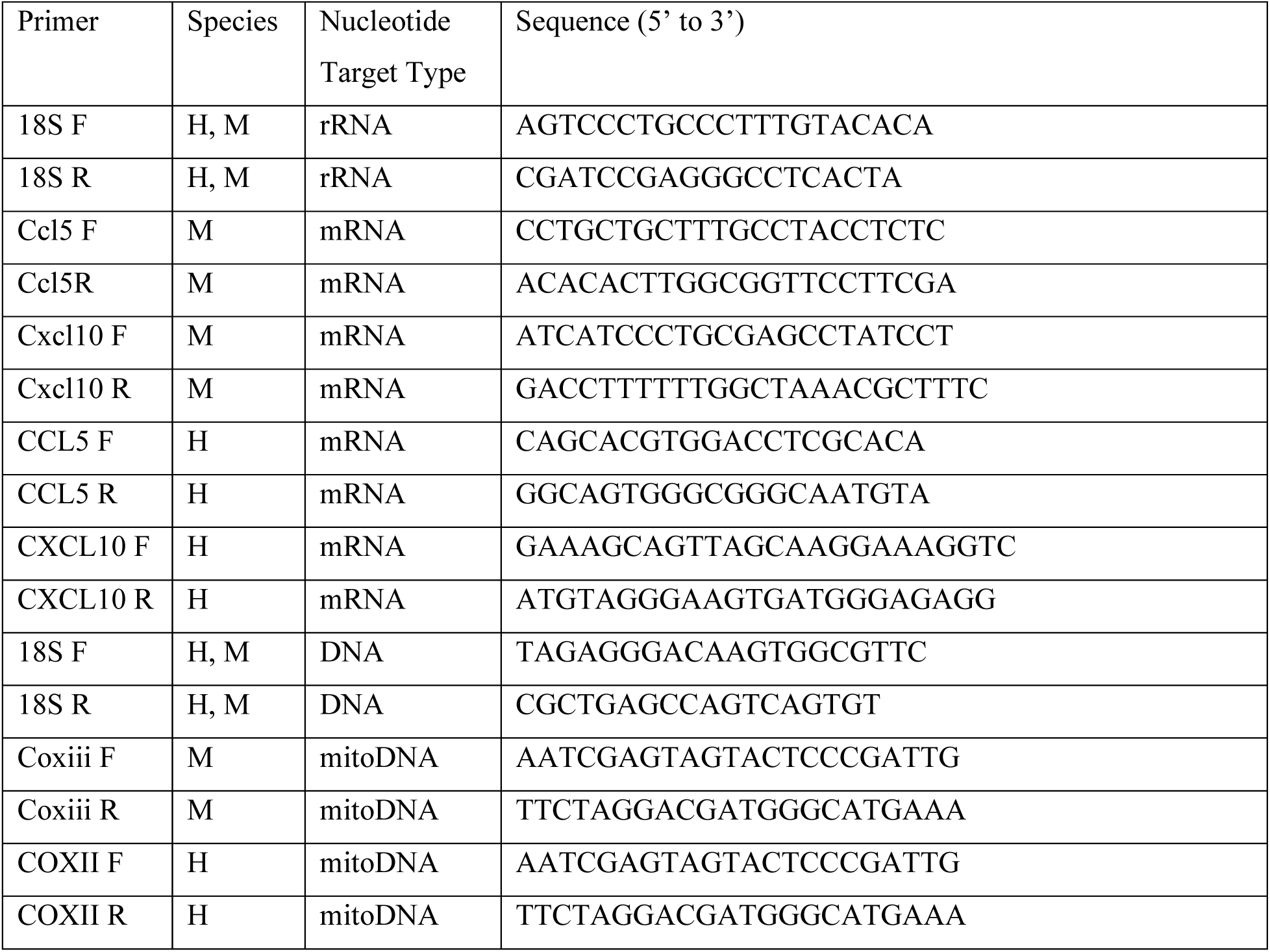
Primer Information.

**Figure S1.**
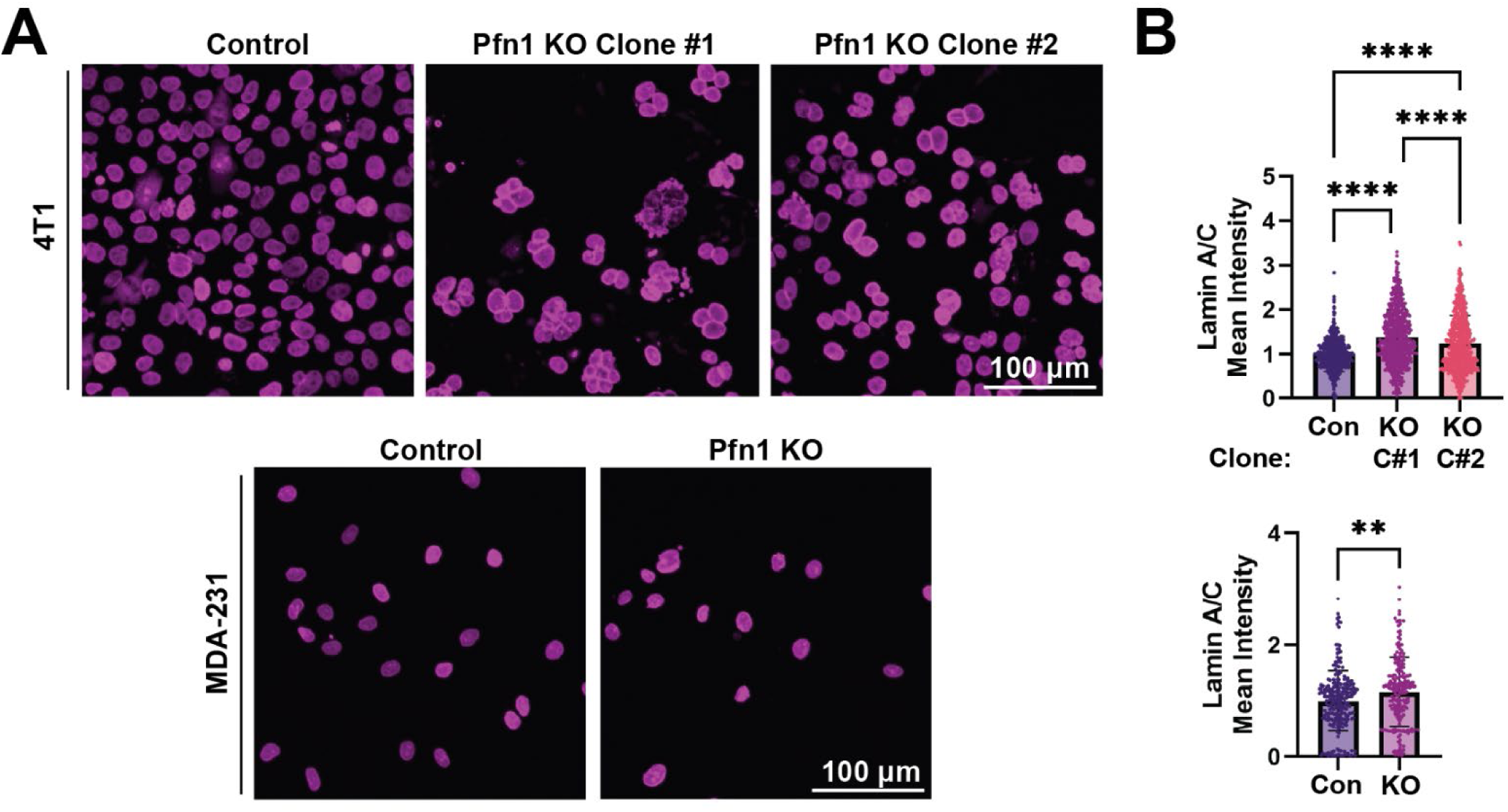
Pfn1 KO results in an increase in nuclear envelope Lamin-A/C levels. **A)** Representative immunofluorescence images of Lamin-A/C-stained control vs Pfn1 KO 4T1 and MDA-231 cells. **B)** Quantification of the mean intensity of Lamin-A/C staining in the nuclear lamina region. Data pooled from three independent experiments. A student’s T test or One-way ANOVA was used to compare between groups. (4T1: Con = 678 cells, Pfn1 KO (C#1) = 743 cells, Pfn1 KO (C#2) =740 cells; MDA-231: Con = 239 cells, Pfn1 KO = 207 cells). ** p < 0.01, **** p < 0.0001

**Figure S2.**
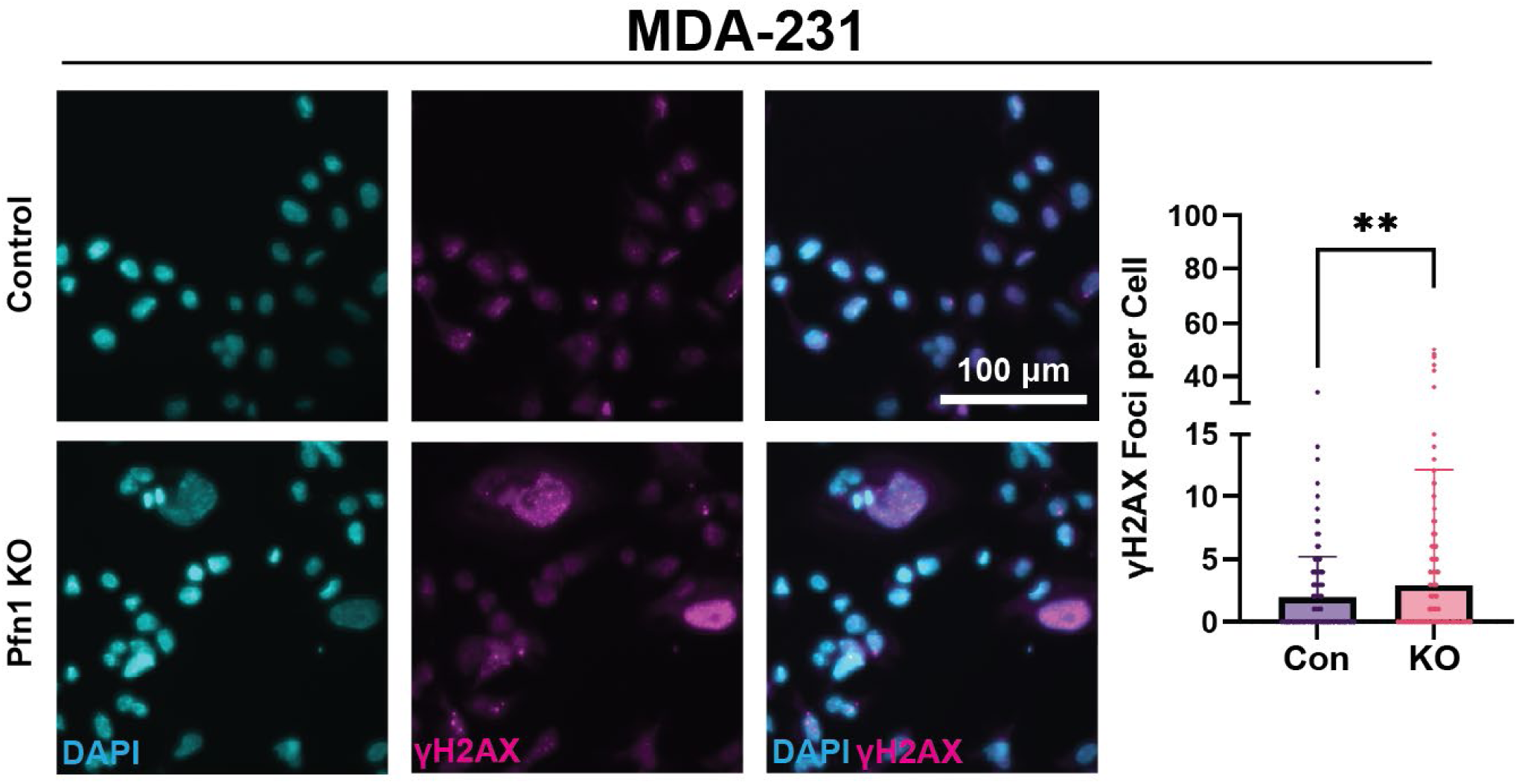
Pfn1 KO results in an increase in the number of γH2AX foci in MDA-231 cells. Representative Images and quantification of the number of γH2AX foci in the nuclei of Control and Pfn1 KO cells. Data pooled from three independent experiments. Con = 1695 cells, Pfn1 KO = 954 cells. A student’s test was used to compare between groups. ** p < 0.01

**Figure S3.**
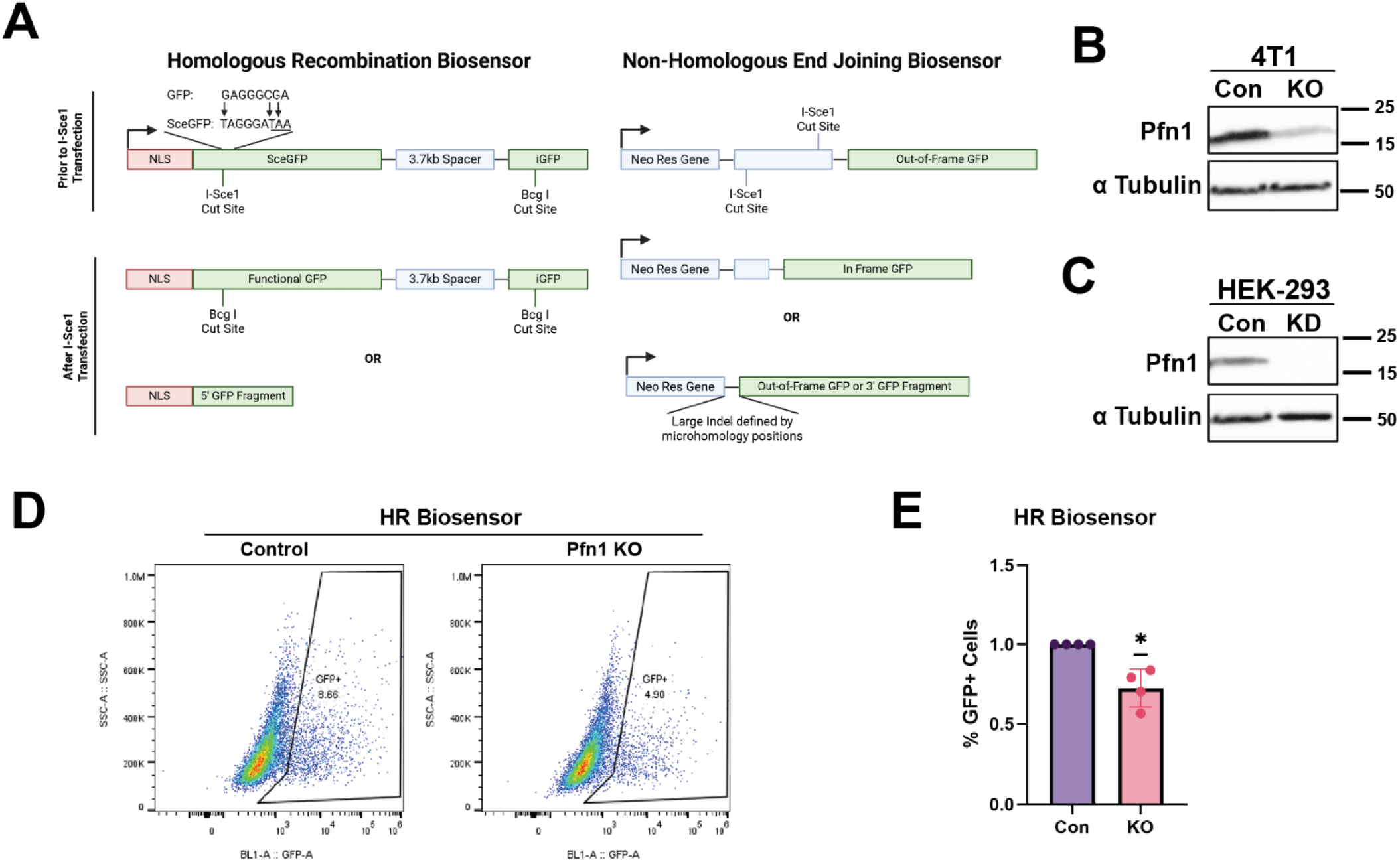
Additional validation of reduced DSB repair proficiency of Pfn1-deficient cells. **A)** Schematics demonstrating the principle of the HR and NHEJ biosensor assays. **B)** Western blot demonstrating cas9-mediated KO of Pfn1 in a non-GFP expressing 4T1 cells used for the HR and NHEJ biosensor experiments (tubulin blot serves as the loading control). **C)** Immunoblots demonstrating siRNA-mediated Pfn1 knockdown in HEK-293 cells (tubulin blot serves as the loading control). **D)** Representative flow cytometry data showing the relative proportion of HR-proficient control and Pfn1 KO 4T1 cells as measured by GFP^+^ cells. **E)** Relative proportion of HR-proficient control and Pfn1 knockdown HEK-293 cells (data summarized from four independent experiments). A one-sample T-test was used to compare experimental values to a hypothetical mean of 1.

**Figure S4.**
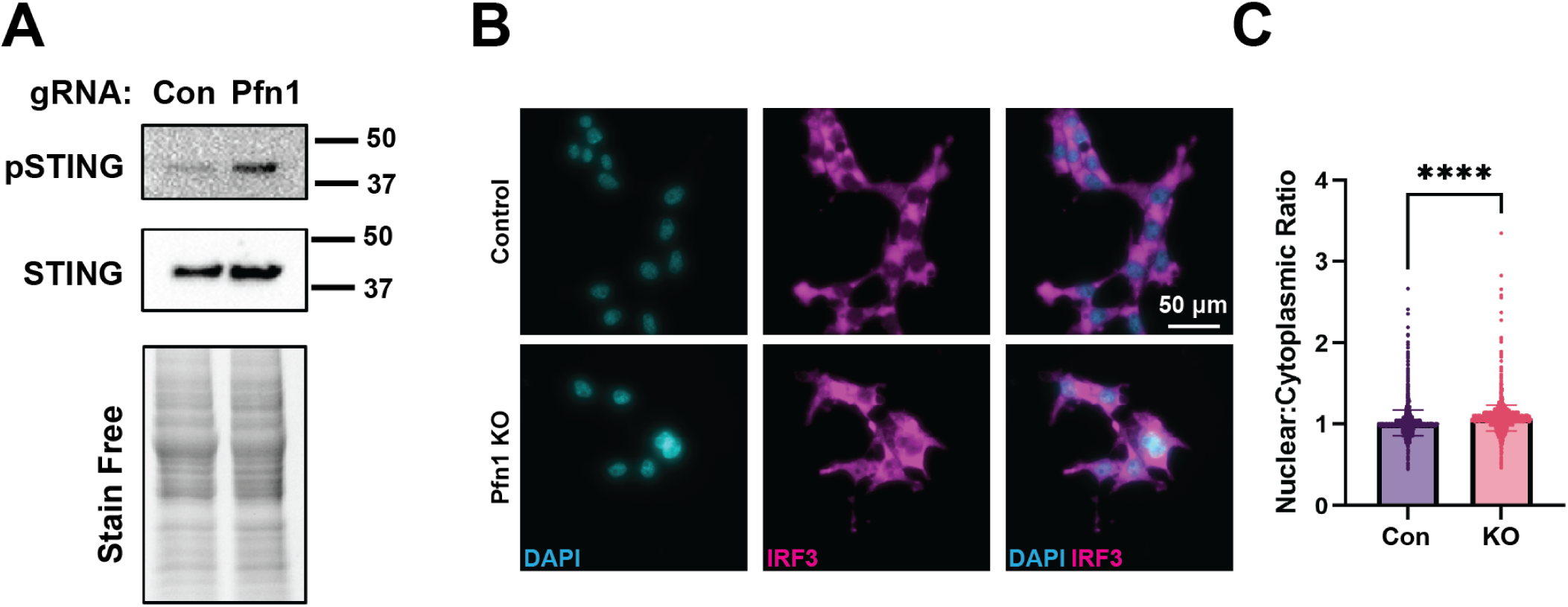
STING pathway activation is elevated in Pfn1 KO 4T1 cells. **A)** Immunoblot of STING and pSTING (Ser365) confirming elevated activation of STING in Polyclonal Pfn1 KO 4T1 cells expressing a second Pfn1-targetting gRNA (a different gRNA than the one used in the rest of the studies). **B)** Representative immunofluorescence images of IRF3 stained control and Pfn1 KO cells. **C)** Realtive comparison between the mean nuclear-to-cytoplasmic ratio of the staining intensity of IRF3 between the two groups. Data pooled from three independent experiments. Con = 4203 cells, Pfn1 KO (C#2) = 2915 cells. A student’s T-test was used to compare between groups.

**Figure S5.**
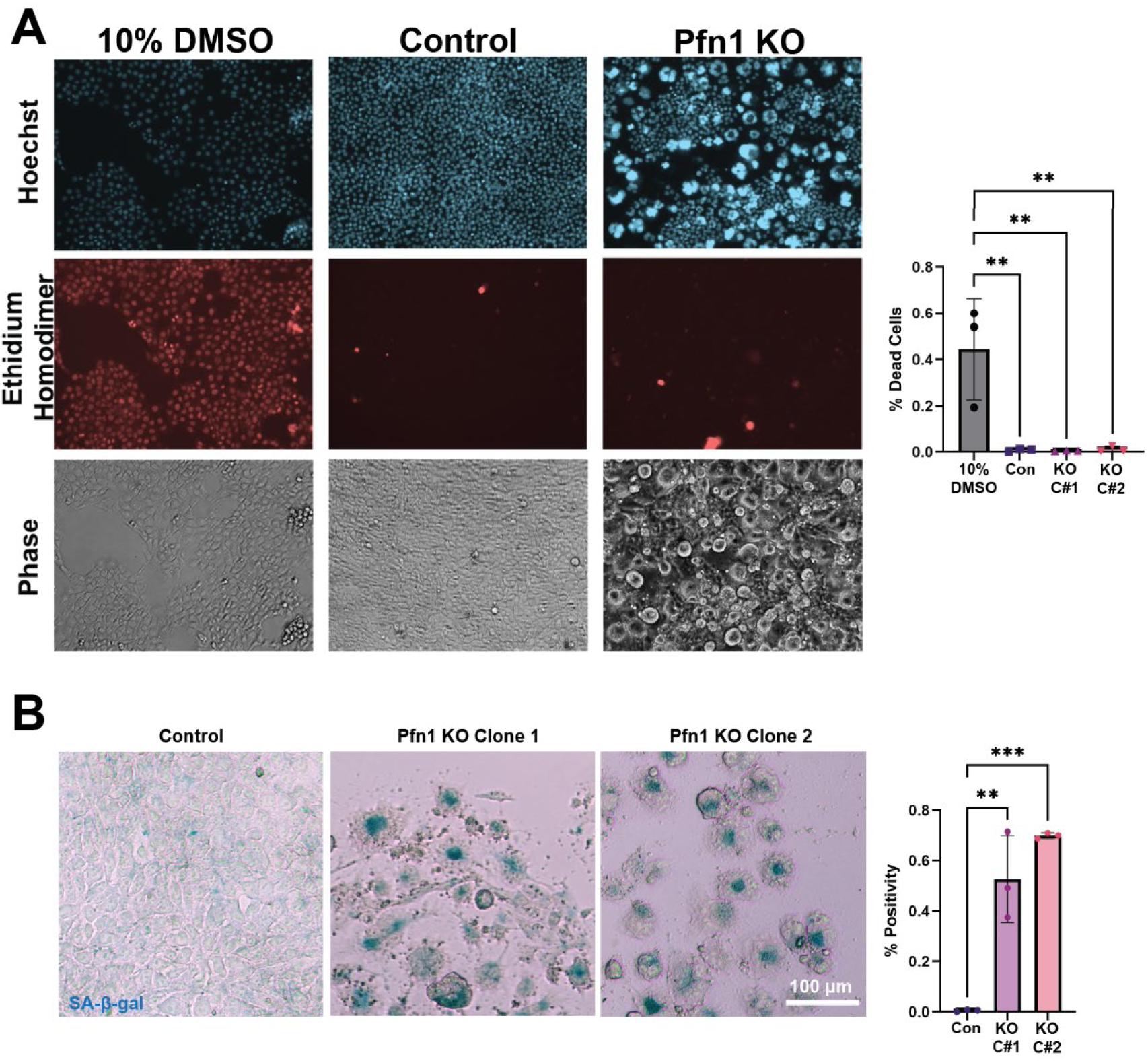
Pfn1 KO induces senescence but does not compromise viability of 4T1 cells. **A)** Representative phase contrast and fluorescence images of live cells stained with Hoechst and Ethidium homodimer and associated quantification. 10% DMSO treatment for 2h represents a positive control for the dead cell quantification. **B)** Representative color bright field images of X-gal based SA-β-gal staining and associated quantification. For both quantifications, a one-way ANOVA was used to compare the percent of total cells with ethidium homodimer or SA-β-gal staining. Each data point represents the average percentage of SA-β-gal positive cells (acquired from multiple fields) in three independent experiments.

**Figure S6.**
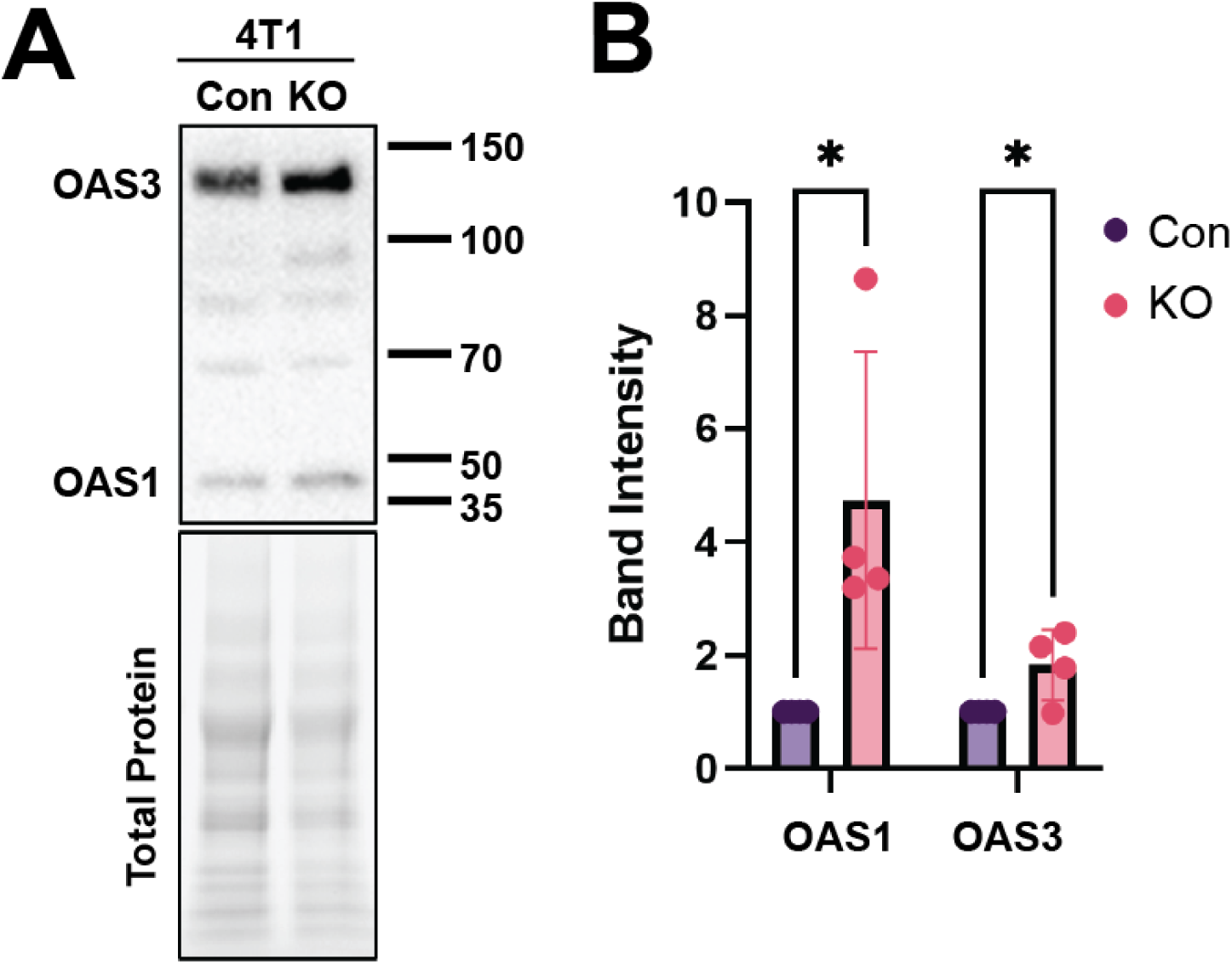
Expression of OAS1/3, a feature of a Type 1 Interferon response, is upregulated in Pfn1 KO cells. **A, B**) Representative immunoblots and the associated quantification showing the relative expression levels of OAS1/3 in the indicated sublines of 4T1. A one-sample T-test was used to compare experimental values to a hypothetical mean of 1. The Pfn1 KO group for 4T1 cells represents clone C#2.

**Figure S7.**
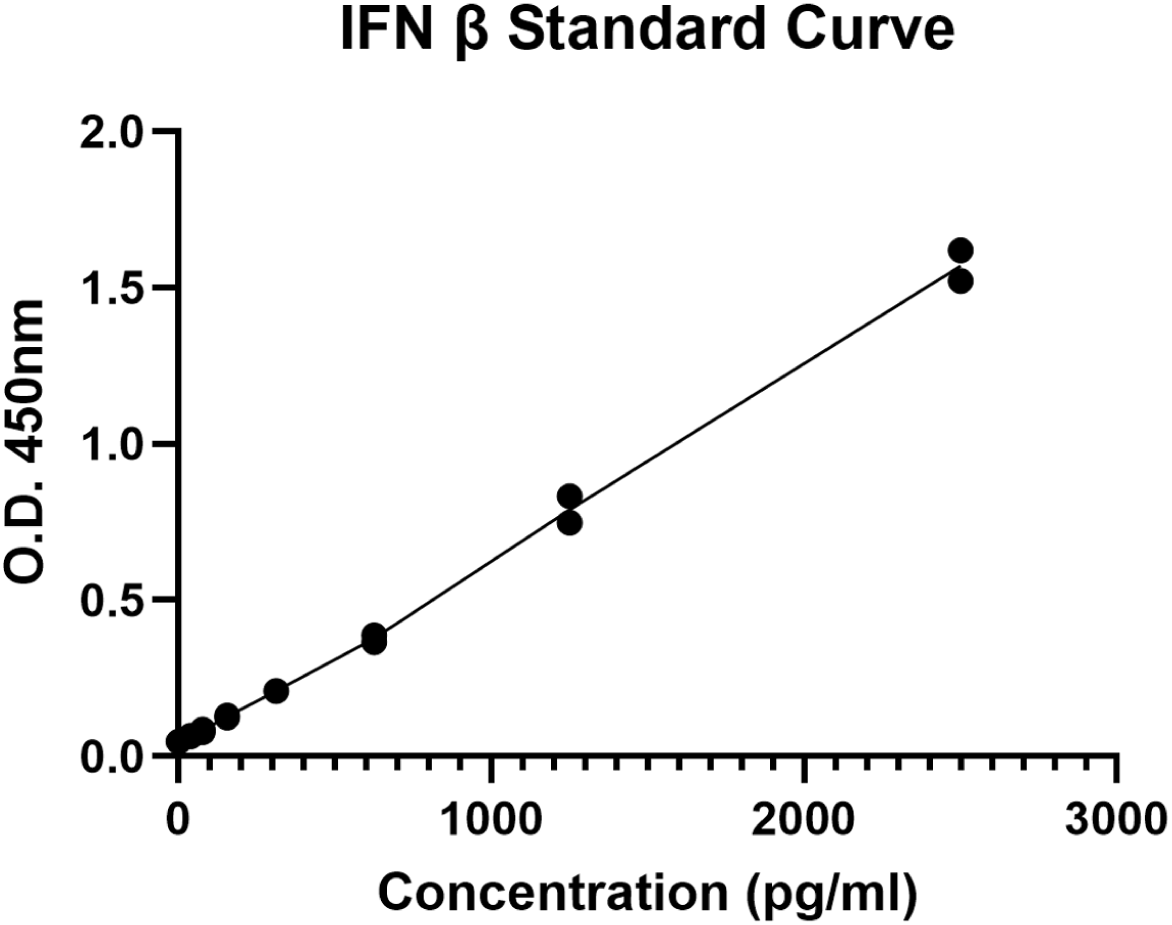
Standard curve for the IFN-β ELISA quantification.

**Figure S8.**
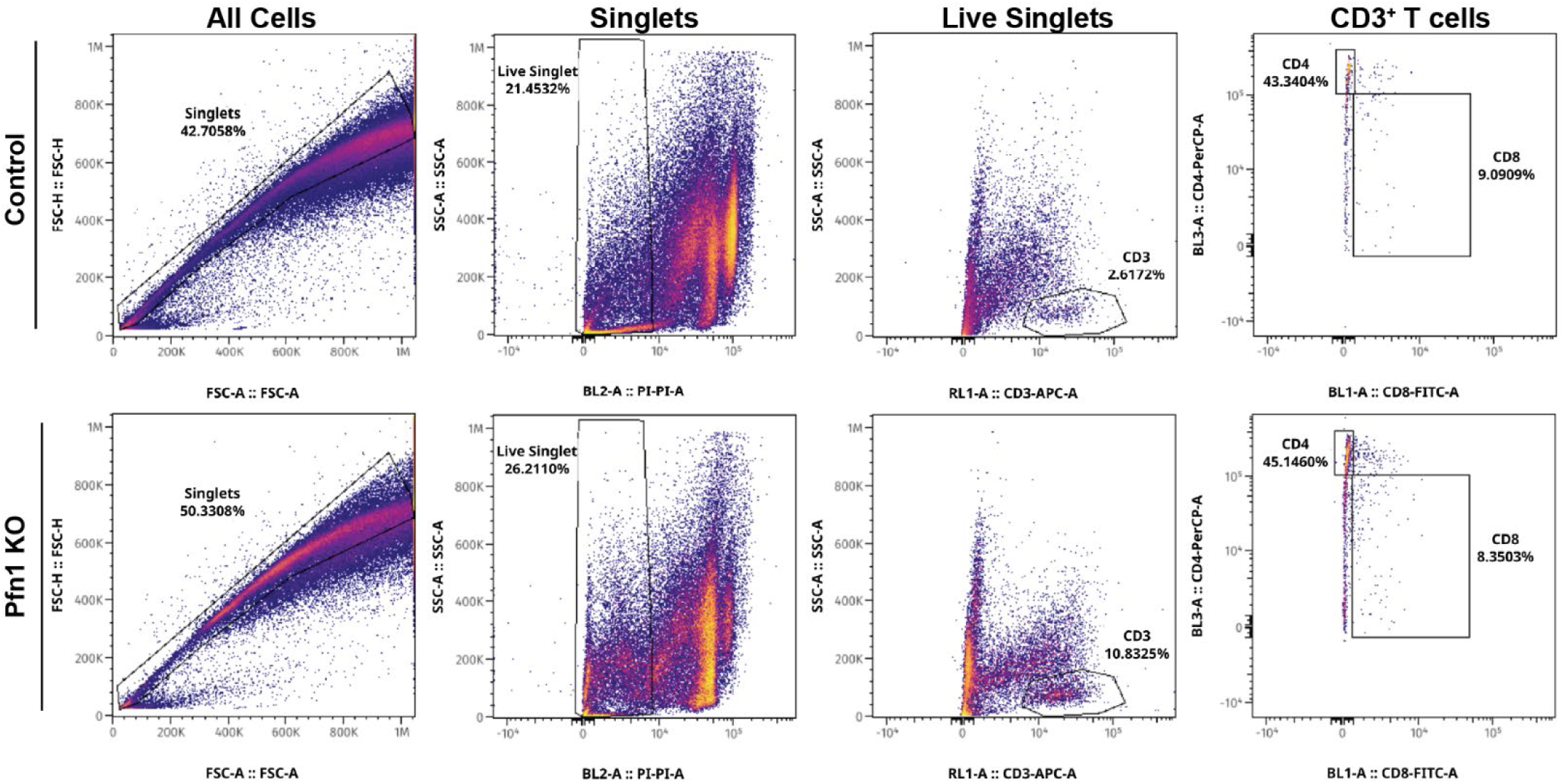
Depiction of immune cell population flow cytometric gating strategy. Representative flow cytometry bivariate plots demonstrating the gating strategy employed for immune cell population estimation in tumors harvested 14 days after inoculation (i.e., 7 days after dox induction). The Pfn1 KO group for 4T1 cells represents clone C#2.

**Figure S9.**
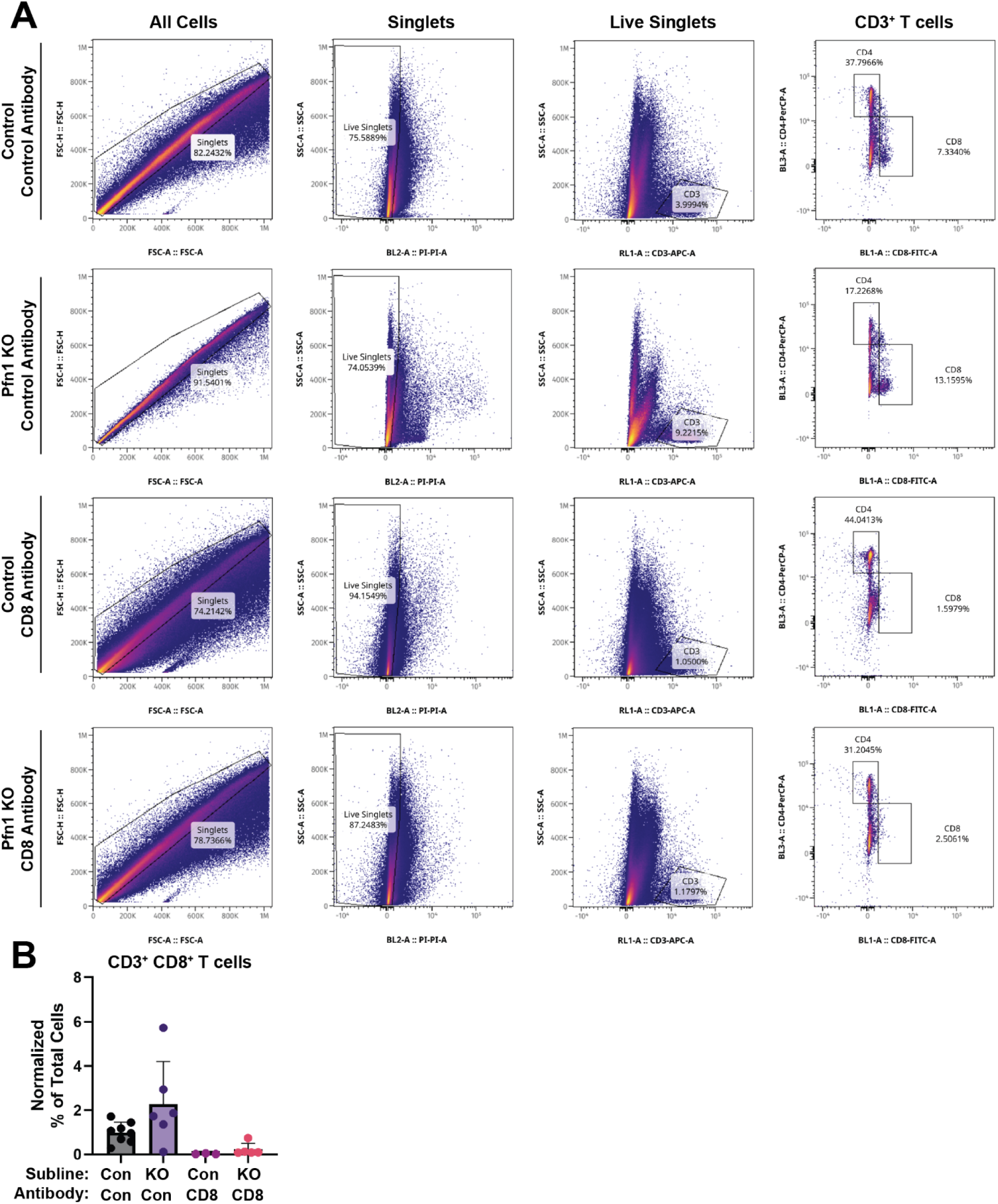
Validation of CD8^+^ T cell depletion in mice. **A)** Representative flow cytometry bivariate plots demonstrating the gating strategy employed for immune cell population estimation in tumors with or without CD8^+^ T cell depletion harvested 17 days after inoculation (i.e., 10 days after dox induction). **B)** Quantification of the percentage of CD3^+^, CD8^+^ live single cells. The Pfn1 KO group represents clone C#2.

## Notes

### Competing Interest Statement

The authors have declared no competing interest.

## REFERENCES

1. Z. Ding, Y. H. Bae, P. Roy, Molecular insights on context-specific role of profilin-1 in cell migration. Cell adhesion & migration 6, 442–449 (2012).

2. Z. Ding, D. Gau, B. Deasy, A. Wells, P. Roy, Both actin and polyproline interactions of profilin-1 are required for migration, invasion and capillary morphogenesis of vascular endothelial cells. Experimental cell research 315, 2963–2973 (2009).

3. B. A. Cisterna et al., Prolonged depletion of profilin 1 or F-actin causes an adaptive response in microtubules. The Journal of cell biology 223 (2024).

4. Z. Ding, A. Lambrechts, M. Parepally, P. Roy, Silencing profilin-1 inhibits endothelial cell proliferation, migration and cord morphogenesis. Journal of cell science 119, 4127–4137 (2006).

5. C. Jiang et al., A balanced level of profilin-1 promotes stemness and tumor-initiating potential of breast cancer cells. Cell Cycle 16, 2366–2373 (2017).

6. R. T. Böttcher et al., Profilin 1 is required for abscission during late cytokinesis of chondrocytes. The EMBO journal 28, 1157–1169 (2009).

7. J. L. Henty-Ridilla, M. A. Juanes, B. L. Goode, Profilin Directly Promotes Microtubule Growth through Residues Mutated in Amyotrophic Lateral Sclerosis. Curr Biol 27, 3535–3543.e3534 (2017).

8. M. M. C. Ricci et al., Actin-binding protein profilin1 is an important determinant of cellular phosphoinositide control. J Biol Chem 300, 105583 (2024).

9. A. Orenberg et al., Molecular insights into Profilin1-dependent regulation of cellular phosphatidylinositol-(4,5)-bisphosphate. Journal of cell science 10.1242/jcs.265025 (2026).

10. T. A. Read et al., The actin binding protein profilin 1 localizes inside mitochondria and is critical for their function. EMBO reports 25, 3240–3262 (2024).

11. Z. Ding et al., Profilin-1 downregulation has contrasting effects on early vs late steps of breast cancer metastasis. Oncogene 33, 2065–2074 (2014).

12. W. Yao et al., Profilin-1 suppresses tumorigenicity in pancreatic cancer through regulation of the SIRT3-HIF1α axis. Molecular cancer 13, 187 (2014).

13. N. Wu et al., Profilin 1 obtained by proteomic analysis in all-trans retinoic acid-treated hepatocarcinoma cell lines is involved in inhibition of cell proliferation and migration. Proteomics 6, 6095–6106 (2006).

14. K. Shen et al., Guttiferone K suppresses cell motility and metastasis of hepatocellular carcinoma by restoring aberrantly reduced profilin 1. Oncotarget 7, 56650–56663 (2016).

15. Z. Wang, Z. Shi, L. Zhang, H. Zhang, Y. Zhang, Profilin 1, negatively regulated by microRNA-19a-3p, serves as a tumor suppressor in human hepatocellular carcinoma. Pathol Res Pract 215, 499–505 (2019).

16. Y. J. Cheng et al., Silencing profilin-1 inhibits gastric cancer progression via integrin β1/focal adhesion kinase pathway modulation. World journal of gastroenterology 21, 2323–2335 (2015).

17. Y. Wang et al., Profilin 1 Induces Tumor Metastasis by Promoting Microvesicle Secretion Through the ROCK 1/p-MLC Pathway in Non-Small Cell Lung Cancer. Front Pharmacol 13, 890891 (2022).

18. A. Allen et al., Actin-binding protein profilin1 promotes aggressiveness of clear-cell renal cell carcinoma cells. J Biol Chem 295, 15636–15649 (2020).

19. J. R. Karamchandani et al., Profilin-1 expression is associated with high grade and stage and decreased disease-free survival in renal cell carcinoma. Hum Pathol 46, 673–680 (2015).

20. F. Scotto di Carlo, L. Pazzaglia, T. Esposito, F. Gianfrancesco, The Loss of Profilin 1 Causes Early Onset Paget’s Disease of Bone. J Bone Miner Res 35, 1387–1398 (2020).

21. Z. Wei et al., Mutations in Profilin 1 Cause Early-Onset Paget’s Disease of Bone With Giant Cell Tumors. J Bone Miner Res 36, 1088–1103 (2021).

22. J. Janke et al., Suppression of tumorigenicity in breast cancer cells by the microfilament protein profilin 1. The Journal of experimental medicine 191, 1675–1686 (2000).

23. L. Zou et al., Profilin-1 is a negative regulator of mammary carcinoma aggressiveness. Br J Cancer 97, 1361–1371 (2007).

24. C. J. Lee, M. J. Yoon, D. H. Kim, T. U. Kim, Y. J. Kang, Profilin-1; a novel regulator of DNA damage response and repair machinery in keratinocytes. Mol Biol Rep 48, 1439–1452 (2021).

25. X. Tian et al., Profilin1 is required for prevention of mitotic catastrophe in murine and human glomerular diseases. The Journal of clinical investigation 133 (2023).

26. C. Rim et al., Nuclear Profilin-1 for DNA Damage Repair Is Involved in Phagocytic Impairment of Senescent Microglia. Glia 73, 1707–1726 (2025).

27. K. J. Mackenzie et al., cGAS surveillance of micronuclei links genome instability to innate immunity. Nature 548, 461–465 (2017).

28. M. Requesens, F. Foijer, H. W. Nijman, M. de Bruyn, Genomic instability as a driver and suppressor of anti-tumor immunity. Front Immunol 15, 1462496 (2024).

29. E. M. Hatch, A. H. Fischer, T. J. Deerinck, M. W. Hetzer, Catastrophic nuclear envelope collapse in cancer cell micronuclei. Cell 154, 47–60 (2013).

30. M. Fenech et al., Molecular mechanisms of micronucleus, nucleoplasmic bridge and nuclear bud formation in mammalian and human cells. Mutagenesis 26, 125–132 (2011).

31. H. Duan et al., Micronuclei: origins, assays, mechanisms, diseases and treatments. Signal Transduct Target Ther 11 (2026).

32. Y. Kalukula, A. D. Stephens, J. Lammerding, S. Gabriele, Mechanics and functional consequences of nuclear deformations. Nature reviews. Molecular cell biology 23, 583–602 (2022).

33. E. P. Rogakou, D. R. Pilch, A. H. Orr, V. S. Ivanova, W. M. Bonner, DNA double-stranded breaks induce histone H2AX phosphorylation on serine 139. J Biol Chem 273, 5858–5868 (1998).

34. R. Ceccaldi, B. Rondinelli, A. D. D’Andrea, Repair Pathway Choices and Consequences at the Double-Strand Break. Trends in cell biology 26, 52–64 (2016).

35. A. J. Pierce, R. D. Johnson, L. H. Thompson, M. Jasin, XRCC3 promotes homology-directed repair of DNA damage in mammalian cells. Genes & development 13, 2633–2638 (1999).

36. N. Bennardo, A. Cheng, N. Huang, J. M. Stark, Alternative-NHEJ is a mechanistically distinct pathway of mammalian chromosome break repair. PLoS Genet 4, e1000110 (2008).

37. T. Costelloe et al., The yeast Fun30 and human SMARCAD1 chromatin remodellers promote DNA end resection. Nature 489, 581–584 (2012).

38. M. Fournier et al., KAT2-mediated acetylation switches the mode of PALB2 chromatin association to safeguard genome integrity. eLife 11 (2022).

39. S. H. Lee et al., The SET domain protein Metnase mediates foreign DNA integration and links integration to nonhomologous end-joining repair. Proceedings of the National Academy of Sciences of the United States of America 102, 18075–18080 (2005).

40. R. Broderick et al., EXD2 promotes homologous recombination by facilitating DNA end resection. Nature cell biology 18, 271–280 (2016).

41. A. A. Sartori et al., Human CtIP promotes DNA end resection. Nature 450, 509–514 (2007).

42. Z. Zhu, W. H. Chung, E. Y. Shim, S. E. Lee, G. Ira, Sgs1 helicase and two nucleases Dna2 and Exo1 resect DNA double-strand break ends. Cell 134, 981–994 (2008).

43. T. Sugiyama, S. C. Kowalczykowski, Rad52 protein associates with replication protein A (RPA)-single-stranded DNA to accelerate Rad51-mediated displacement of RPA and presynaptic complex formation. J Biol Chem 277, 31663–31672 (2002).

44. S. M. Harding et al., Mitotic progression following DNA damage enables pattern recognition within micronuclei. Nature 548, 466–470 (2017).

45. L. Sun, J. Wu, F. Du, X. Chen, Z. J. Chen, Cyclic GMP-AMP synthase is a cytosolic DNA sensor that activates the type I interferon pathway. Science 339, 786–791 (2013).

46. Z. Tong, J. P. Zou, S. Y. Wang, W. W. Luo, Y. Y. Wang, Activation of the cGAS-STING-IRF3 Axis by Type I and II Interferons Contributes to Host Defense. Adv Sci (Weinh) 11, e2308890 (2024).

47. Y. Li et al., Activation of RNase L is dependent on OAS3 expression during infection with diverse human viruses. Proceedings of the National Academy of Sciences of the United States of America 113, 2241–2246 (2016).

48. S. S. Withers et al., Effect of stimulator of interferon genes (STING) signaling on radiation-induced chemokine expression in human osteosarcoma cells. PloS one 18, e0284645 (2023).

49. Z. Hong et al., STING inhibitors target the cyclic dinucleotide binding pocket. Proceedings of the National Academy of Sciences of the United States of America 118 (2021).

50. W. Witke, J. D. Sutherland, A. Sharpe, M. Arai, D. J. Kwiatkowski, Profilin I is essential for cell survival and cell division in early mouse development. Proceedings of the National Academy of Sciences of the United States of America 98, 3832–3836 (2001).

51. B. Schrörs et al., Multi-Omics Characterization of the 4T1 Murine Mammary Gland Tumor Model. Frontiers in oncology 10, 1195 (2020).

52. L. Hui, Y. Zheng, Y. Yan, J. Bargonetti, D. A. Foster, Mutant p53 in MDA-MB-231 breast cancer cells is stabilized by elevated phospholipase D activity and contributes to survival signals generated by phospholipase D. Oncogene 25, 7305–7310 (2006).

53. T. Stüven, E. Hartmann, D. Görlich, Exportin 6: a novel nuclear export receptor that is specific for profilin.actin complexes. The EMBO journal 22, 5928–5940 (2003).

54. B. R. Schrank et al., Nuclear ARP2/3 drives DNA break clustering for homology-directed repair. Nature 559, 61–66 (2018).

55. B. J. Belin, T. Lee, R. D. Mullins, DNA damage induces nuclear actin filament assembly by Formin -2 and Spire-½ that promotes efficient DNA repair. [corrected]. eLife 4, e07735 (2015).

56. C. Zhu et al., Profilin-1 regulates DNA replication forks in a context-dependent fashion by interacting with SNF2H and BOD1L. Nature communications 13, 6531 (2022).

57. Z. Zhao et al., Loss of Nuclear Profilin 1 Triggers Oncogenic Reprogramming of Mammary Epithelial Cells Through Dysregulated DNA Replication in Breast Cancer. J Breast Cancer 28, 333–346 (2025).

58. U. S. Srinivas, B. W. Q. Tan, B. A. Vellayappan, A. D. Jeyasekharan, ROS and the DNA damage response in cancer. Redox Biol 25, 101084 (2019).

59. A. Giampetruzzi et al., Modulation of actin polymerization affects nucleocytoplasmic transport in multiple forms of amyotrophic lateral sclerosis. Nature communications 10, 3827 (2019).

60. C. H. Wu et al., Mutations in the profilin 1 gene cause familial amyotrophic lateral sclerosis. Nature 488, 499–503 (2012).

61. A. Freund, R. M. Laberge, M. Demaria, J. Campisi, Lamin B1 loss is a senescence-associated biomarker. Molecular biology of the cell 23, 2066–2075 (2012).

62. A. Barascu et al., Oxidative stress induces an ATM-independent senescence pathway through p38 MAPK-mediated lamin B1 accumulation. The EMBO journal 31, 1080–1094 (2012).

63. X. Wang et al., ROS/p38MAPK-induced lamin B1 accumulation promotes chronic kidney disease-associated vascular smooth muscle cells senescence. Biochemical and biophysical research communications 531, 187–194 (2020).

64. Z. Zhou et al., Sensing of cytoplasmic chromatin by cGAS activates innate immune response in SARS-CoV-2 infection. Signal Transduct Target Ther 6, 382 (2021).

65. Y. Jing et al., STING couples with PI3K to regulate actin reorganization during BCR activation. Sci Adv 6, eaax9455 (2020).

66. T. Araki, J. G. Williams, Perturbations of the actin cytoskeleton activate a Dictyostelium STAT signalling pathway. Eur J Cell Biol 91, 420–425 (2012).

67. H. Guo et al., Cucurbitacin I inhibits STAT3, but enhances STAT1 signaling in human cancer cells in vitro through disrupting actin filaments. Acta Pharmacol Sin 39, 425–437 (2018).

68. M. Smid et al., Patterns and incidence of chromosomal instability and their prognostic relevance in breast cancer subtypes. Breast cancer research and treatment 128, 23–30 (2011).

69. S. F. Bakhoum et al., Chromosomal instability drives metastasis through a cytosolic DNA response. Nature 553, 467–472 (2018).

70. M. Robson et al., Olaparib for Metastatic Breast Cancer in Patients with a Germline BRCA Mutation. The New England journal of medicine 377, 523–533 (2017).

71. M. Joy, D. Gau, N. Castellucci, R. Prywes, P. Roy, The myocardin-related transcription factor MKL co-regulates the cellular levels of two profilin isoforms. J Biol Chem 292, 11777–11791 (2017).

72. J. Li et al., Co-inhibitory Molecule B7 Superfamily Member 1 Expressed by Tumor-Infiltrating Myeloid Cells Induces Dysfunction of Anti-tumor CD8(+) T Cells. Immunity 48, 773–786.e775 (2018).

73. N. Arnoult et al., Regulation of DNA repair pathway choice in S and G2 phases by the NHEJ inhibitor CYREN. Nature 549, 548–552 (2017).

74. C. Richardson, M. E. Moynahan, M. Jasin, Double-strand break repair by interchromosomal recombination: suppression of chromosomal translocations. Genes & development 12, 3831–3842 (1998).

75. A. S. Jahun, F. Sorgeloos, I. G. Goodfellow, An optimized protocol for the extraction and quantification of cytosolic DNA in mammalian cells. STAR Protoc 5, 102913 (2024).

76. D. Aran, M. Sirota, A. J. Butte, Systematic pan-cancer analysis of tumour purity. Nature communications 6, 8971 (2015).

77. M. Menzel et al., Accurate tumor purity determination is critical for the analysis of homologous recombination deficiency (HRD). Transl Oncol 35, 101706 (2023).

78. D. W. Yao, N. G. Balanis, E. Eskin, T. G. Graeber, A linear mixed model approach to gene expression-tumor aneuploidy association studies. Scientific reports 9, 11944 (2019).

